# Molecular profiling of high-level athlete skeletal muscle after acute exercise – a systems biology approach

**DOI:** 10.1101/2023.04.11.535671

**Authors:** Stefan M Reitzner, Eric B Emanuelsson, Muhammad Arif, Bogumil Kaczkowski, Andrew TJ Kwon, Adil Mardinoglu, Erik Arner, Mark A. Chapman, Carl Johan Sundberg

## Abstract

Life-long high-level exercise training leads to improvements in physical performance and multi-tissue adaptation following changes in molecular pathways. While skeletal muscle baseline differences between exercise-trained and untrained individuals have been previously investigated, it remains unclear how acute exercise multi-omics are influenced by training history. We recruited and extensively characterized 24 individuals categorized as endurance athletes, strength athletes or control subjects. Multi-omics profiling was performed from skeletal muscle before and at three time-points after endurance or resistance exercise sessions. Timeseries multi-omics analysis revealed distinct differences in molecular processes such as fatty- and amino acid metabolism and for transcription factors such as HIF1A and the MYF-family between both exercise history and acute form of exercise. Furthermore, we found a “transcriptional specialization effect” by transcriptional narrowing and intensification. Finally, we performed multi-omics network analysis and clustering, providing a novel resource of skeletal muscle transcriptomic and metabolomic profiling in highly trained and untrained individuals.

## INTRODUCTION

High-level physical training results in exercise modality-specific adaptations in skeletal muscle that are partly due to transient as well as accumulating changes in gene expression and metabolite levels ^1–3^. Following each exercise bout, their levels are influenced by an interaction between signaling molecules and the transcriptional machinery, over time contributing to the gradual remodeling of skeletal muscle^3^. The structural and biochemical remodeling that has accrued in trained skeletal muscle, in turn, influences its transcriptional and metabolic response to acute exercise bouts^4, 5^. Previous investigations of skeletal muscle gene expression following acute exercise bouts or exercise training programs^4, 6–8^ have been limited either by exercise intervention duration (weeks to months) or training status of the population studied (untrained to recreationally active). Other investigations of acute exercise effects have focused on blood only^9^. Accordingly, to overcome such limitations, we recruited two groups of long-term trained (15+ years) endurance and resistance athletes and an age-matched untrained control group. All participants performed acute bouts of endurance and resistance exercise in a cross-over design before and after which biopsies were obtained from skeletal muscle on which multi-omics analyses were conducted.

The aim was to identify key molecular pathways connected to physical performance capabilities, comparing subject groups and modality of acute exercise to better understand the molecular trajectories of acute exercise. During subject characterization, several performance parameters were measured. The association between such markers and molecular level pathways and metabolites can be of great value for the identification of processes that drive skeletal muscle health and performance. Increased understanding of the molecular characteristics in life-long trained athletes could provide valuable knowledge for training protocol optimization by refocusing on training modalities that trigger key performance-relevant adaptations such as for example introducing endurance or resistance elements in SG or EG athletes respectively. Further, such increased understanding can be clinically relevant for the development of strategies to prevent, treat and assess the status of diseases like diabetes, metabolic syndrome, cardiovascular disease, cancer or enabling beneficial exercise-induced effects when exercise is not an option. While efforts have previously been made to understand and identify such processes in different tissues or with short duration exercise interventions^9^, no study to date has systematically investigated the influence of life-long high-level training on the transcriptome, metabolome and transcription factor motifs of skeletal muscle in humans. Finally, by comparing the intra-individual response to two different forms of acute exercise, a deeper understanding of common or distinctively different adaptational responses to endurance and resistance exercise can be facilitated by thorough molecular profiling.

Results from our comprehensive characterization of the transcriptomic, metabolomic and regulatory response to acute exercise show extensive differences based on training history, amongst others due to sports-specific specializations in energy metabolic pathways. We reveal distinct molecular differences between acute endurance and resistance exercise and, by using correlation analysis identify key metabolic processes that could act as key drivers to performance output such as carnitine and amino acid metabolisms. Taken together, we deliver for the first time, a multi-omics characterization of skeletal muscle from highly trained individuals. Additionally, we provide a valuable, easy to use public resource (http://www.inetmodels.com/ (Multi-omics network → Study-specific Networks → crossEX (Reitzner et al. 2023)) for future investigations of the molecular effects of acute exercise and long-term adaptation and highlight the potential of multi-omics profiling for the investigation of training effects.

## RESULTS

### Subject Characterization and Analyses Performed

Highly characterized healthy male subjects (n=24) between 33 and 52 were recruited into three groups based on self-reported training history and a series of physiological performance tests (V̇O2peak, 1 repetition maximum and peak torque knee extension, Table 1): life-long high-level trained endurance (EG) and strength (SG) athletes and age-matched untrained healthy participants (CG). Inclusion criteria were based on leg strength, VO_2_peak test and training history and identical to our previously published cross-sectional study^10^. Included research subjects underwent a series of skeletal muscle biopsies before and after two acute exercise interventions separated by 1-2 months as depicted in Figure 1A. Subsequently, RNA sequencing (transcriptome), liquid and gas chromatography-mass spectrometry (metabolome, lipidome), and *in-silico* genome scale metabolic modelling (GEM) and transcription factor motif activity response analysis (MARA) were performed (Figure 1B). In addition to 11 physiological and histological parameters (see Table 1), the final dataset included a total of 20223 analytes, composed of: 15701 genes, 279 metabolites, 4164 reporter metabolites - markers of in-silico inferred metabolic flux, and 79 transcription factor motifs (Figure 1C).

**Figure 1:**
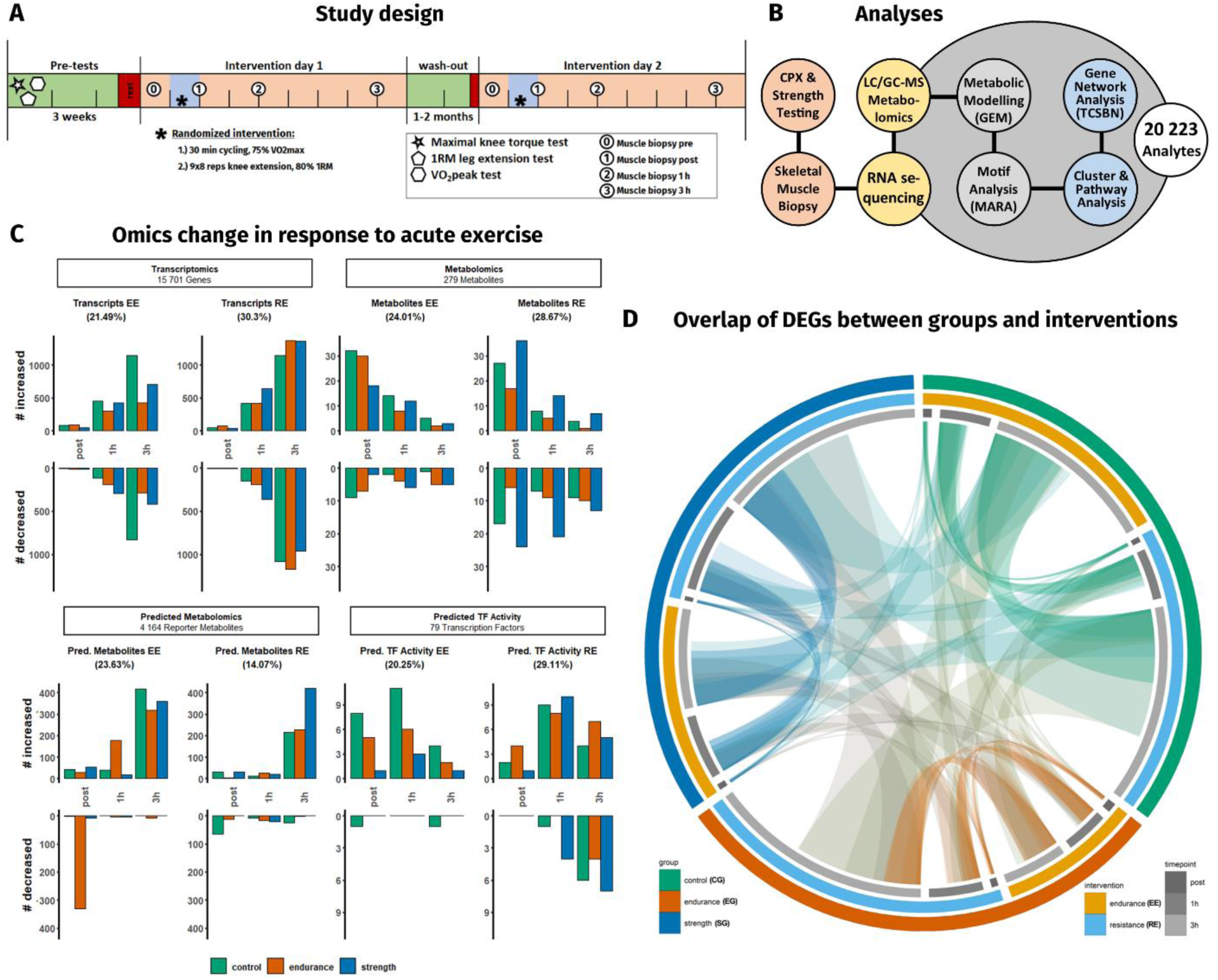
Study Design, Molecular Response to Exercise and overlap of DEGs. (A) Overview over the study design. Two bouts of endurance (EE) or resistance exercise (RE) were performed in a randomized cross-over design and skeletal muscle biopsies taken at 4 timepoints. (B) Cardiopulmonary (CPX) and strength testing and analyses performed. (C) Multi-omics response to acute exercise by subject group with the percentage of total significantly affected by EE or RE. (D) Overlap of differentially expressed genes compared to the respective pre timepoint. Widths of arches and segment lengths are proportional to the number of DEGs.

**Table 1:** Basic characterization of research subjects. Values are group mean ± SD. (*: p < 0.05 compared to both the other groups; ##: p < 0.05 compared to CG

| group | control (CG) | endurance (EG) | strength (SG) |
| --- | --- | --- | --- |
| n | 8 | 8 | 8 |
| age (yrs) | 44 ± 6 | 42 ± 5 | 39 ± 6 |
| height (cm) | 181 ± 6 | 182 ± 4 | 180 ± 5 |
| weight (kg) | 76 ± 8 | 73 ± 4 | 92 ± 12 * |
| BMI (kg/m<sup>2</sup>) | 23.1 ± 2.4 | 22.3 ± 0.5 | 28.3 ± 2.9 * |
| leg strength (Nm) | 180.6 ± 30.8 | 198.3 ± 25.5 | 286.4 ± 28.1 * |
| $\dot{V}O_{2\text{max}}$ (ml·kg<sup>-1</sup>·min<sup>-1</sup>) | 36.2 ± 4.4 | 69 ± 7.2 * | 40.2 ± 6.9 |
| $\dot{V}E/\dot{V}CO_2$ slope | 35.8 ± 5.9 * | 28.9 ± 2.8 | 30.2 ± 1.6 |
| CS activity (nmol·min<sup>-1</sup>·mg protein<sup>-1</sup>) | 5.0 ± 0.5 | 10.1 ± 1.5 * | 4.2 ± 0.7 |
| type 1 fibers | 44.6 % ± 4.4 | 61.4 % ± 3.9 * | 43 % ± 3.2 |
| type 2 fibers | 55.4 % ± 4.4 | 38.6 % ± 3.9 * | 57 % ± 3.2 |
| type 1 CSA (μm<sup>2</sup>) | 4841.7 ± 561.2 * | 6048.3 ± 257.8 | 6660.6 ± 522.3 |
| type 2 CSA (μm<sup>2</sup>) | 5369.5 ± 860.3 | 6632 ± 665.2 # | 9184 ± 415.2 * |
| nuclei/type 1 fiber | 2.6 ± 0.6 | 3.4 ± 0.5 # | 4.4 ± 0.6 * |
| nuclei/type 2 fiber | 2.9 ± 0.9 | 2.6 ± 0.8 | 4.5 ± 0.9 * |

### Baseline physiological, biochemical and histological differences

By design, leg strength was significantly higher in the strength group (SG) compared to both control group (CG) and endurance group (EG; Table 1). V̇O2peak in EG was significantly higher than both other groups and SG leg strength significantly higher than both EG and CG. Similarly, skeletal muscle citrate synthase (CS) activity was significantly higher in EG than both other groups, a consequence of long-term endurance training^11^. Skeletal muscle type 1 fiber proportion was significantly higher in EG compared with both other groups (Table 1, Figure S1), a difference that has been observed before^12, 13^. Muscle fiber cross-sectional area (CSA) was significantly lower in CG for type 1 compared to both other groups. EG and SG had a similar type 1 CSA but a significantly larger type 2 CSA was observed in SG (Table 1, Figure S1).

### Molecular inter- and intra-group differences at baseline

#### Variability of omics

Descriptive statistical analyses on our dataset (Figure S2A,B, Figure S2C-F) demonstrated that within-group coefficient of variation (CoV) of analytes at baseline was similar in all groups but varied depending on omics-level (Figure S2A left), demonstrating a consistent intra-group variability of omics across groups at baseline with no significant group difference. Previous publications have reported long-term exercise training to have a consolidating effect for individual molecular analytes^14^ or in response to acute exercise^10^. Our results showed that intra-group analyte-wise variation was similar across groups regardless of training status (Figure S2A, Figure S2F), suggesting that the molecular adaptation of high-level athletes lies in the fine-tuning of pathways and individual genes and metabolites rather than in the overall omics dimension. When comparing intra-individual gene expression between the two pre timepoints, small differences were found, with a mean fold change close to zero, and only 3.12-3.96% of all transcripts displaying a fold change of 0.5 or higher (Figure S2C). These results provide an interesting benchmark of inherent, day-to-day shift of individual baseline gene expression. These can be the product of slight differences in daily conditions regarding nutrition, sleep, immune system status or other factors. While we are confident that we tightly controlled for the some of the above-mentioned factors insofar possible, remaining variation can be a product of random or environmental factors such as seasonality, which up to 10% of all genes in skeletal muscle might be influenced by^15^. However with the average distance between intervention days being 55 days this should not be a major factor. Taken together, our data indicates a low level of baseline gene expression variability.

#### Gene expression and metabolomic group differences

Comparing gene expression of groups at baseline, results showed smaller differences between CG and SG (57 DEGs) than between the EG and both other groups (1363 with CG and 919 with SG; Supplement A, Table S1), similar to what was previously shown by us^10^. Using gene set enrichment analysis (GSEA), we show that these DEGs are associated with mitochondrial pathways enriched in the endurance group, while differences between CG and SG are mainly attributed to immune system and structural terms (Figure S2D).

On metabolomic level, SG showed to have high inter-individual diversity of metabolite levels of sorbitol, gluconic acid and erythritol (Figure S2B), all previously reported to play a major role in skeletal muscle glucose metabolism in rodents^16–18^. We found metabolic baseline differences between groups (see Supplement A, Table S2 and Figure S2E). Specifically, carnitines were more abundant in the EG compared to both other groups, representing about 20% of the significantly different metabolites. Carnitines act as intracellular chaperone molecules for beta-oxidation via the carnitine-palmitoyltransferase system which have a rate-limiting role^19, 20^ for the transport of fatty acids across the mitochondrial membranes. They have been reported to be elevated in circulating blood of high-level trail runners^21^. High concentrations of different length fatty acids bound to their carnitine transporters, such as propryonyl- or butyrylcarnitine can be interpreted both as a higher potential for beta-oxidative capacity and higher levels of ongoing beta-oxidation at rest^22^. Such an adaptation is of great benefit for endurance exercise, but generally of less relevance to resistance exercise. About 30% of all metabolites that were significantly more abundant in SG than in CG were the phospholipid groups of phosphatidylcholines (PC) and phosphatidylethanolamines (PE), which have previously been described as relevant for membrane fluidity^23^, insulin receptor- and membrane protein dynamics^24, 25^ and mitochondrial structure^26^ (Figure S2E). The positive effect of RE on insulin sensitivity and membrane fluidity has previously been demonstrated^27, 28^, and has been shown to be a consequence of a more flexible recruitment of GLUT4 into the plasma membrane^29^, thus aiding glucose uptake. Together, both these metabolic PC-PE and carnitine profiles found in SG and EG, are consistent with the respective functional requirements of high-level endurance and resistance exercise.

### Molecular Response to Acute Exercise

The acute response to exercise has been suggested to follow a time-specific choreography, modulated by acute exercise modality and training background^9, 30^. To better understand this modulation, we compared trajectories of differentially expressed genes (DEGs). An overview can be found in Figure S3A. Together with the higher number of DEGs in response to RE (Figure 1C), this data shows a more robust activation of *M. vastus lateralis* gene expression in response to RE compared to EE. Exercise modality showed to have a bigger role for gene activation than training background as indicated by stronger connections between same type acute exercise than within subject group (Figure 1D, Figure S3B). The most unique set of genes showed to be at the 3h timepoint with as much as 84% of genes in EG performing RE (Figure S3A). However, compared to the 3h timepoint, the earlier timepoints resulted in a more coordinated gene expression response as measured by numbers of highly enriched pathways identified by GSEA. Such a pattern might be explained by EG being highly capable of meeting acute energy requirements of RE, but exhibiting a chaotic, less coordinated response, similar to untrained individuals at later stages. In contrast, as a first indicator of a group difference in the response to acute exercise, CG and SG show a high degree of significantly enriched pathways at the 3h timepoint (Figure 2A), particularly in response to RE rather than EE.

**Figure 2:**
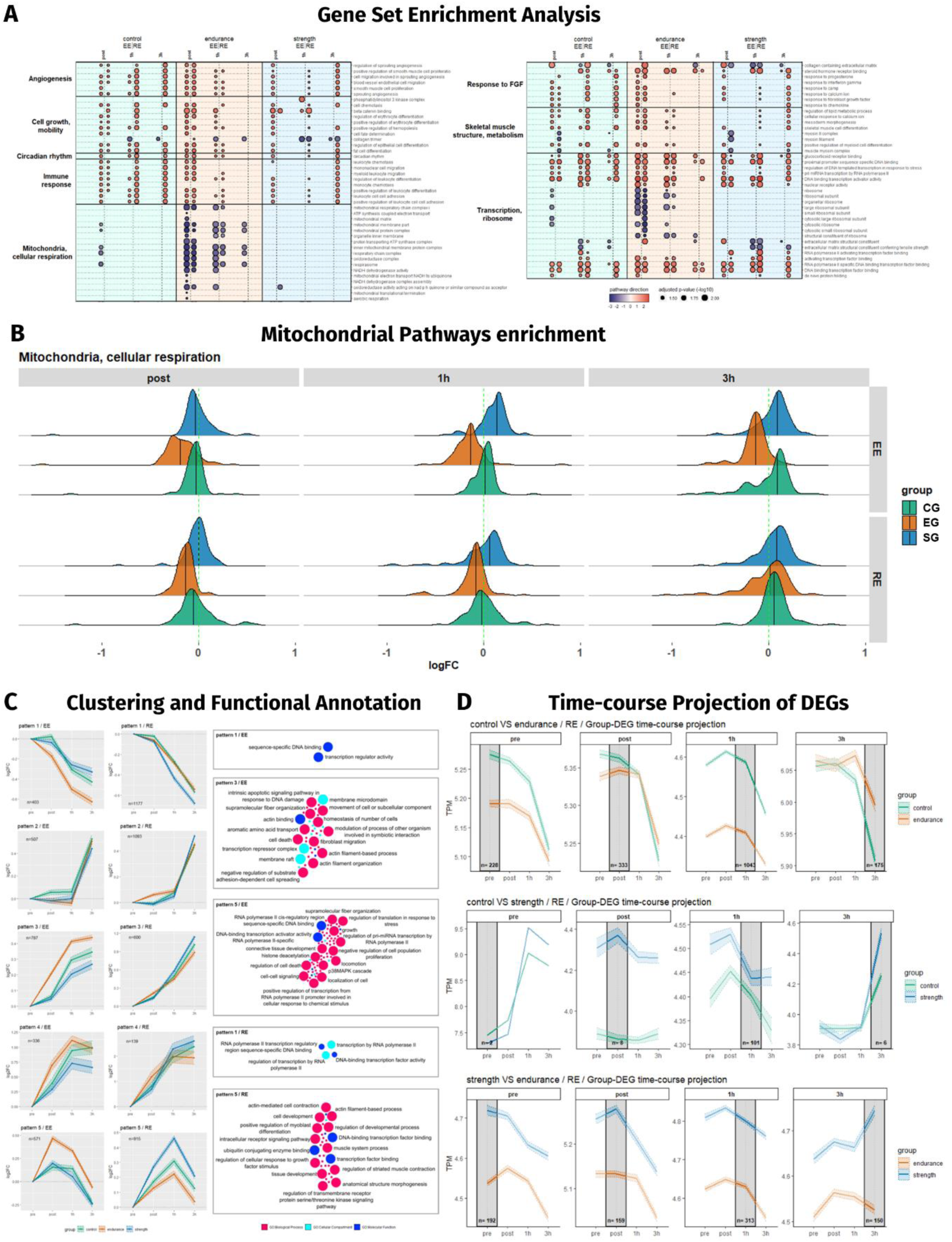
Functional analysis of the transcriptomic response to acute endurance and resistance exercise. (A) Gene set enrichment analysis (GSEA) of DEGs for acute endurance (EE) and resistance exercise (RE). Individual pathways are summarized by supergroups. (B) Ridge plot of mitochondria and cellular respiration pathways molecular signatures from A. (C) Gene cluster identity from unsupervised clustering of genes of the endurance group performing EE (EE patterns, left) and the strength group performing RE (RE patterns, right) are used to compare the same sets of genes in all groups. Corresponding GSEA is shown on the right. (D) Projection (plotting of DEGs from one timepoint across the remaining timepoints; origin comparison timepoint with grey background) of DEGs comparing groups at individual timepoints to the remaining timepoints (mean ± confidence interval).

#### Functional annotation of differential gene expression after acute exercise

Further GSEA analysis showed angiogenesis-, immune system-, growth factor- and transcription-related pathways to be upregulated across all groups and interventions (Figure 2A). However, pathways related to mitochondria/cellular respiration and transcription/ribosome were uniquely downregulated in EG in response to both EE and RE (Figure 2A), particularly at the post timepoint (Figure 2B). Mitochondria are a major target of skeletal muscle adaptation, particularly following endurance training, influencing their biogenesis^31^, plasticity^32^, increasing cristae density^32^ and potential citrate cycle throughput as measured by CS activity^33–35^, which was significantly higher in EG compared to both CG and SG (Table 1). As shown above, EG unlike both CG and SG has increased baseline enrichment of pathways contributing to mitochondrial biogenesis and building up their function (Figure S2D). The aforementioned downregulation (Figure 2A) might represent this high mitochondrial capacity of EG, providing sufficient amounts of energy substrates during acute exercise, and immediate downregulation upon cessation of exercise beyond baseline levels due to a complete saturation with energy substrates. Both, EG and SG are highly specialized in their respective disciplines, suggesting their acute exercise transcriptomic response is highly optimized. Considering this, we were interested in how the sets of genes stimulated in highly adapted athletes respond in both the groups naïve to that form of exercise, CG and SG being EE-naïve, and CG and EG being SG-naïve. To do so, we performed unsupervised clustering of DEGs of EG performing EE, and SG performing RE. The same genes were then plotted as clusters for all groups, enabling a direct group comparison of each cluster of DEGs. Results show the same general shape, thus similar transcriptomic programs being stimulated by acute exercise regardless of training history. However, following EE, a more pronounced response in EG (pattern 1, 3, 5; Figure 2C) was found, while the same was true in SG following RE (pattern 1, 5; Figure 2C). This suggests the differences between highly trained and exercise-naïve individuals in the omics dimension lie in a fine-tuning of the degree of gene expression, not a large re-arrangement of the entire transcriptome, notwithstanding possible large differences on the level of individual genes. Functional profiling of patterns 1, 3 and 5 in EE showed an association with regulation of transcription, structural, myofibrillar and translation related terms. RE patterns 1 and 5 were associated with regulation of transcription, developmental process, morphogenesis and growth. Overall, less distinct group differences were found following RE, which is in line with the above-mentioned similarity between CG and SG, suggesting a smaller transcriptional adaptation to RE than to EE. We also performed this analysis based on clustering from CG, with some but less striking differences (Figure S3C; pattern 2 EE and 5 RE). Further comparison of group differences additionally supported this previously mentioned notion of a broad but fine-tuning of acute gene regulation (Figure 2D, Figure S3D). For this, we used group-comparison DEGs from each timepoint and plotted summary-level expression across all timepoints (Figure 2D). Across the majority of comparisons in response to both RE and EE, gene trajectories are largely parallel, however staying clearly separated in most cases, especially comparing SG and EG (Figure 2D bottom panel, Figure S3C). Together, this suggests a similar transcriptomic expression pattern, however at a different expression level, meaning that the acute form of exercise dictates the transcriptomic response, irrespective of training background, which rather influences expression amplitude.

#### Generation of a genome-scale metabolic network

To expand the perspective on the dynamics of metabolites in response to acute exercise, we used the transcriptomic data to generate a genome-scale metabolic network (GEM) by integration of all known human metabolic equations and pathways^36^. Most notably, the generated GEM showed a large change in EE stimulated metabolic flux at the post timepoint in the EG but not in CG and SG. EG showed to have 362 significantly affected reporter metabolites, of which 332 were in the negative direction (Figure 1C). CG and SG showed 45 and 62 affected reporter metabolites respectively, mainly in the positive direction. Using KEGG, the EG reporter metabolites were largely mapped to energy-related metabolic processes such as TCA cycle and fatty acid metabolism (Figure S5A). These results further reinforce our GSEA-based findings of a large-scale downregulation of mitochondria and cellular respiration pathways immediately following EE in EG, but not in the CG and SG (Figure 2A,B). This drastic, immediate down-regulation following EE might indicate a tighter control over up- and downregulation of an acute exercise transcriptomic profile in EG, for example an abrupt cessation of transcription of energy-pathway related genes immediately after the end of the acute exercise and a drop in ATP demand, compared to both CG and SG.

#### Acute effect of exercise on the metabolome

After this first predictive perspective on metabolic flux, the abundance of 279 metabolites were directly measured by targeted LC/GC-MS. Following unsupervised *k*-means clustering of significantly changed metabolites^37^, they were annotated by their metabolic superpathways (Figure 3A, Figure S4A). Interestingly, the metabolite sorbitol, a product of aldose reductase and the first step of the polyol pathway, showed to be most responsive to acute exercise in all comparisons. While studies on its effect in skeletal muscle are sparse, some authors have suggested sorbitol can cause hyperosmolar stress, positively affecting glucose uptake^38, 39^. Though not reported to be acutely regulated in human muscle before, its proposed function would be consistent with the energy requirements during and following acute exercise. Cluster analysis further revealed differences in how lipid-associated metabolites respond to acute exercise in EG: Following EE, the majority of lipid-associated metabolites were upregulated (8 metabolites; clusters 3 and 4; Figure 3A), compared to 4 metabolites in response to RE (cluster 3) and 3 out of 4 metabolites in the downregulated cluster being lipid-associated (cluster 5). Closer investigation of the lipid superfamily showed derivates of carnitine oppositely affected in response to EE (4 derivates up-) and RE (3 derivates downregulated). The remaining lipid-associated metabolites were regulated in the same direction (Figure 3B,D). In EG, modification of the carnitine shuttle, trafficking long chain fatty acids into the mitochondrion for beta oxidation, constitutes yet another hallmark of energy flux optimization in skeletal muscle tissue. This process results in a reduction in free carnitine and an increase in carnitine derivates^40^. The ratio between the two has been used as a marker of the utilization of beta-oxidation in metabolic disorders^41, 42^. Similarly, a distinct difference in amino acid-associated metabolites comparing EE and RE in SG was observed (Figure 3A, cluster 4, 5, 6; Figure S4A, cluster 4, 5). A closer investigation revealed a large-scale downregulation in response to RE only (Figure 3C), amongst others the branched chain amino acids (BCAA) valine, leucine and isoleucine, but notably not of ketoleucine, a product and indicator of the ongoing BCAA catabolism. In contrast, the only BCAA downregulated in response to EE was valine (Figure 3D). This response pattern and difference between EE and RE is less clear in EG, with all three BCAAs being downregulated in response to EE (Figure S4C). Together, this indicates a sharper distinction between forms of acute exercise for amino acids (AA) in SG, whereas EG is more optimized for fatty acid trafficking, reflecting exercise background-specific metabolic specializations.

**Figure 3:**
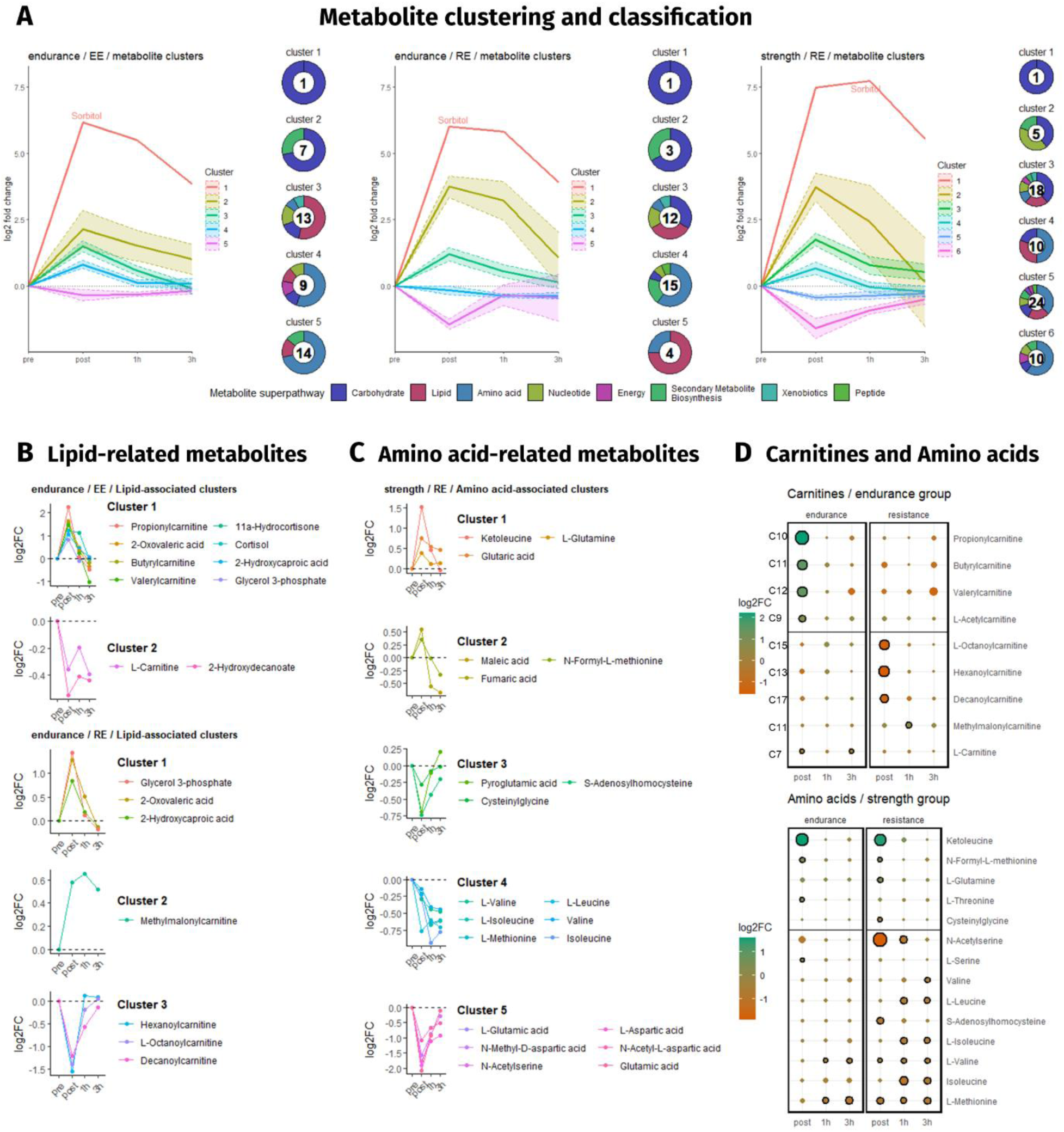
Clustering and classification of metabolomics. (A) Clustering (mean ± confidence interval) and classification of metabolites of endurance (EG) and strength group (SG) in response to acute endurance (EE) and resistance exercise (RE). Numbers in donuts represent the cluster size. (B) Clustering of lipid-associated metabolites in EG in response to EE and RE and (C) of amino acid-associated metabolites in SG in response to RE. (D) Analysis of carnitines in EG and amino acids in SG in response to acute exercise. Size and color of dots represent effect size and direction. Black circles represent statistical significance compared to pre.

#### Transcription factor motif activity analysis

To obtain a better understanding of the regulation of the observed transcriptomic changes, we employed Motif Activity Response Analysis (MARA), which pairs transcriptomics with gene-wise transcription factor motif binding site information^43–45^. By doing so, we were able to discover additional exercise-response dynamics. Unlike DEG which peaks at 3h, most significantly changed transcription factor (TF) motifs peaked at the 1h timepoint, primarily increasing in activity (Supplement A, Table S4). Interestingly, some of these comparisons prominently feature HIF1A and MYFfamily. HIF1A is a regulator of mitochondrial metabolism and hypoxia-related processes^46–48^, with oxygen delivery to skeletal muscle being one of the primary limiting factors of acute endurance performance^49^. HIF1A controls genes responsible for erythropoiesis and angiogenesis, aimed at coping with and resolving the hypoxic condition^50^, but itself relies on a complex regulation involving degradation by the von Hippel-Lindau protein and activation in hypoxic conditions^51, 52^. The MYFfamily motif controls the myogenic regulatory factors MYF5, MYF6, MYOD1 and MYOG^53^. Comparing groups following RE, EG was less likely than SG to activate strength adaptation-associated motifs such as MYFfamily but was, amongst others, more likely to activate the HIF1A motif (Supplement A, Table S4). Surprisingly, while EG exhibited the highest number of DEGs following RE on transcriptomic level, EE induced a higher number of changes on the TF motif level. This difference might be the consequence of a more untargeted, “chaotic” response to an unfamiliar form of exercise as opposed to a targeted activation of functionally connected genes by specific TFs, and could explain both, transcriptomic and TF level effects. As demonstrated by baseline differences, EG is specialized in a markedly different regulatory direction compared to both CG and SG, including a regulatory specialization with a more pronounced stimulation of EE-associated transcription factors, such as HIF1A. This results in a narrower, but more intense stimulation of transcription as indicated above in the functional cluster analysis (Figure 2C). The same logic could also explain the somewhat blunted activity of the MYFfamily motif following RE in EG compared with CG (Supplement A, Table S4). HIF1A and MYFfamily motifs both show an exercise-specific response in a direct comparison of EE and RE (Figure S6A). Importantly, they seem to only be affected in their activity following EE (HIF1A motif) and RE (MYFfamily motif), but not following the other form of exercise (Figure S6B). Furthermore, we show the CREB1 motif, which regulates PPARGC1A, a master regulator of mitochondrial biogenesis^54, 55^, to be stimulated by both EE and RE (Figure S4B).

MARA calculates motif activity based on the expression of all motif-associated genes^45, 56^. To identify the primary genes driving the activity of each motif, we used correlation analysis (“reverse mapping”), revealing a number of candidate genes highly relevant to their respective motif (Figure S6C). Many of these genes have previously been described to play a substantial role in the regulatory response to acute exercise such as HES1 (associated with HIF1A motif), involved in skeletal muscle differentiation^57^, CLIC4 (MYFfamily motif), having a role in angiogenesis^58^, TRIM63, a potential regulator of muscle mass^59^ and CBFB (both CREB1 motif), a negative regulator of MyoD^60^. Using tissue- and cancer-specific biological network analysis (TCSBN) we further confirmed these candidate genes to be functionally connected rather than a random collection of genes (Figure S6D).

### Correlation between baseline physiological and histological features and acute molecular effects

Molecular processes stimulated by acute exercise largely reflect the physical and biochemical requirements of the performance output and the ability of the individual to meet these. To better understand the influence of the individual performance characteristics on skeletal muscle molecular changes at different omics levels following acute exercise, transcripts, metabolites and TFs were correlated with V̇O2peak (EE) and peak knee torque (RE). Between 16 and 30% (mean 21%) of analytes significantly correlated (p<0.05) with V̇O2peak across all timepoints following EE (Figure 4A). In contrast, 0 to 20 % (mean 7%) of analytes correlated with peak leg torque following RE. Endurance training has extensive influence on acute mitochondrial activity and ATP production, and subsequently V̇O2peak. RE, on the other hand, stimulates acute molecular mechanisms that will not immediately reflect long-term adaptation, but rather following hypertrophic processes over time increase peak torque and thus contractile potential. Such a process involves molecular mechanisms such as immune system activation, hypertrophic processes and angiogenesis that take place up to several days following the acute bout^61, 62^. To identify genes highly relevant to EE and RE performance via their proxies V̇O_2_peak and peak torque, acute gene expression was individually summarized as area under the curve (AUC) over time using trapezoid integration. To further deepen the analysis, significantly correlating genes from our study were only kept if they were also found to be significantly regulated by acute exercise in two large meta-analyses by Amar *et al*. and Pillon *et al*.^30, 63^ (Figure 4B). The present analysis showed a stronger correlation of genes following RE (up to r=0.73) than following EE (up to r=-0.58; Figure 4B). Notably, PPARGC1A and myogenesis regulating gene MYF6 in RE were identified in this analysis (Figure 4B). While PPARGC1A has primarily been associated with endurance exercise^64–66^, there is some support for a role in resistance exercise, regulating skeletal muscle hypertrophy^67, 68^. Muscle regeneration and hypertrophy rely on a choreography of myogenic factors to guide a satellite cell from quiescence to differentiation, with MYF6 activity required only during the terminal differentiation process^69^. Its negative correlation can be an indication of this particular MYF6 expression timing. Previous results placed MYF6 activity between 1 and 8 hours following acute exercise^70^, a range included in the present study. Alternatively, this could be explained by a lower stimulation of pro-hypertrophic processes in the already hypertrophy-adapted SG^71^. Next, we used the identified top genes in an attempt to explain inter-individual variance in V̇O2peak and peak torque. Results show that as few as 5 transcripts can explain 41% of the variance of V̇O2peak and 4 transcripts 68% of peak torque variability (Figure 4C). Additionally, to get a better understanding of the processes connected to the identified correlating genes, GSEA was performed for all genes that correlated significantly with peak torque or V̇O2peak. Surprisingly, results show a strong enrichment of TCA-cycle, cellular respiration and ribosome-related terms in connection to the ability to produce high peak torque such as SG. Enrichment of genes correlating with high V̇O2peak was rather focused on receptor binding, transcriptional regulation, the electron transfer chain as well as actin binding (Figure 4D).

**Figure 4:**
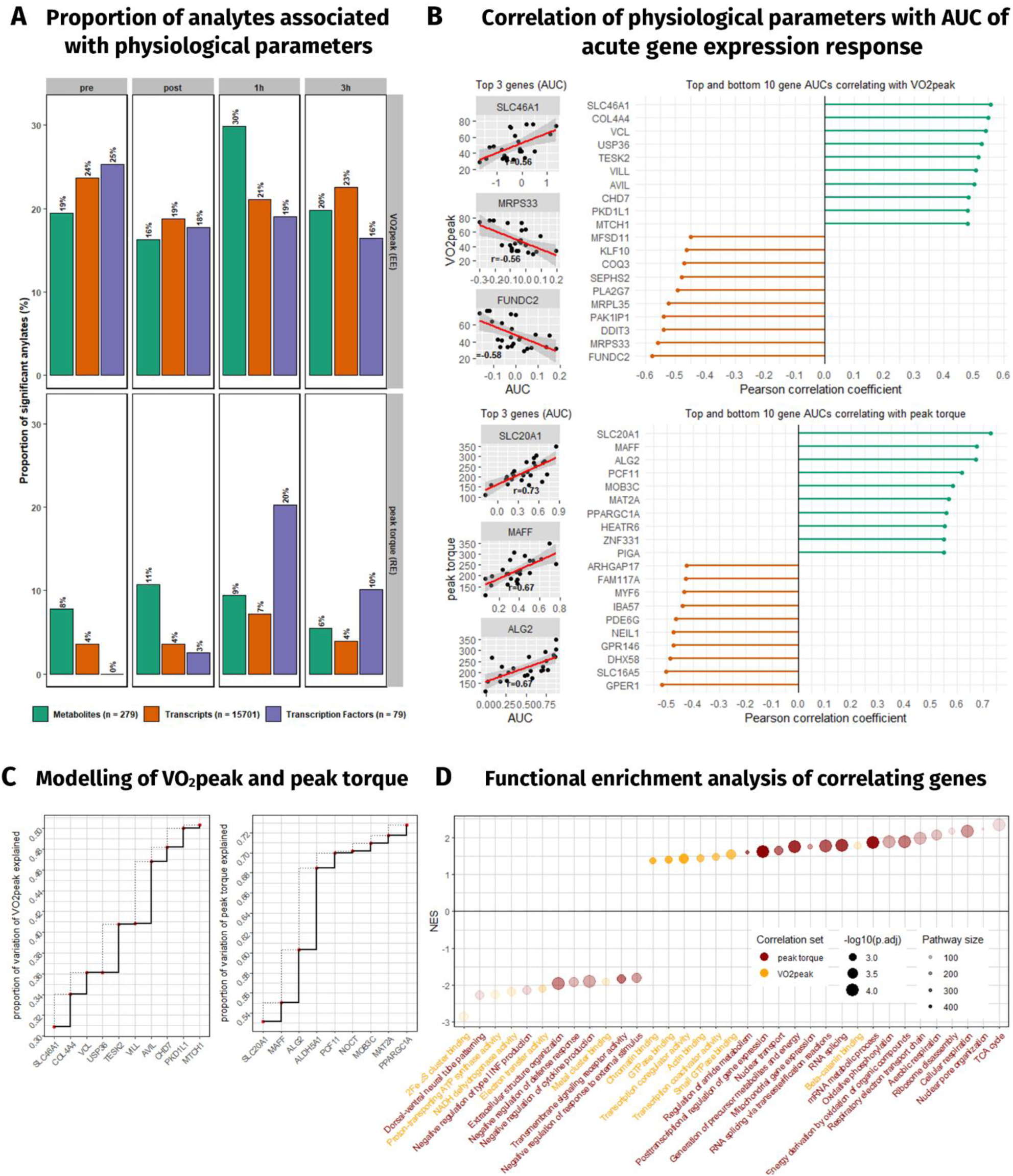
Correlation of physiological parameters with molecular analytes. (A) Proportion of analyte response to EE and RE based on area under curve (AUC) significantly correlating to baseline V̇O2peak and peak torque respectively. Total number of analytes each in legend parenthesis. (B) Top and bottom 10 genes correlating with V̇O2peak and peak torque; top 3 genes shown individually. (C) Modelling of top genes explaining variation of V̇O2peak and peak torque. (D) Gene set enrichment analysis of genes (expressed as normalized enrichment score, NES) sorted by correlation for V̇O2peak and peak torque.

### Multi-omics molecular clustering

To obtain a deeper understanding of the relations between -omics data and individual analytes, transcripts, metabolites, clinical and histological data and predicted TF motifs were used for network integration and cluster analysis at baseline (Figure 5) and in response to acute exercise (Figure 5, Figure 6). This network was then used to determine physiological parameter correlations with individual analytes, for cluster identification and functional annotation. The generated network can be interactively accessed and downloaded for cytoscape via http://www.inetmodels.com/ (Multi-omics network → Study-specific Networks → crossEX (Reitzner et al. 2023).

**Figure 5:**
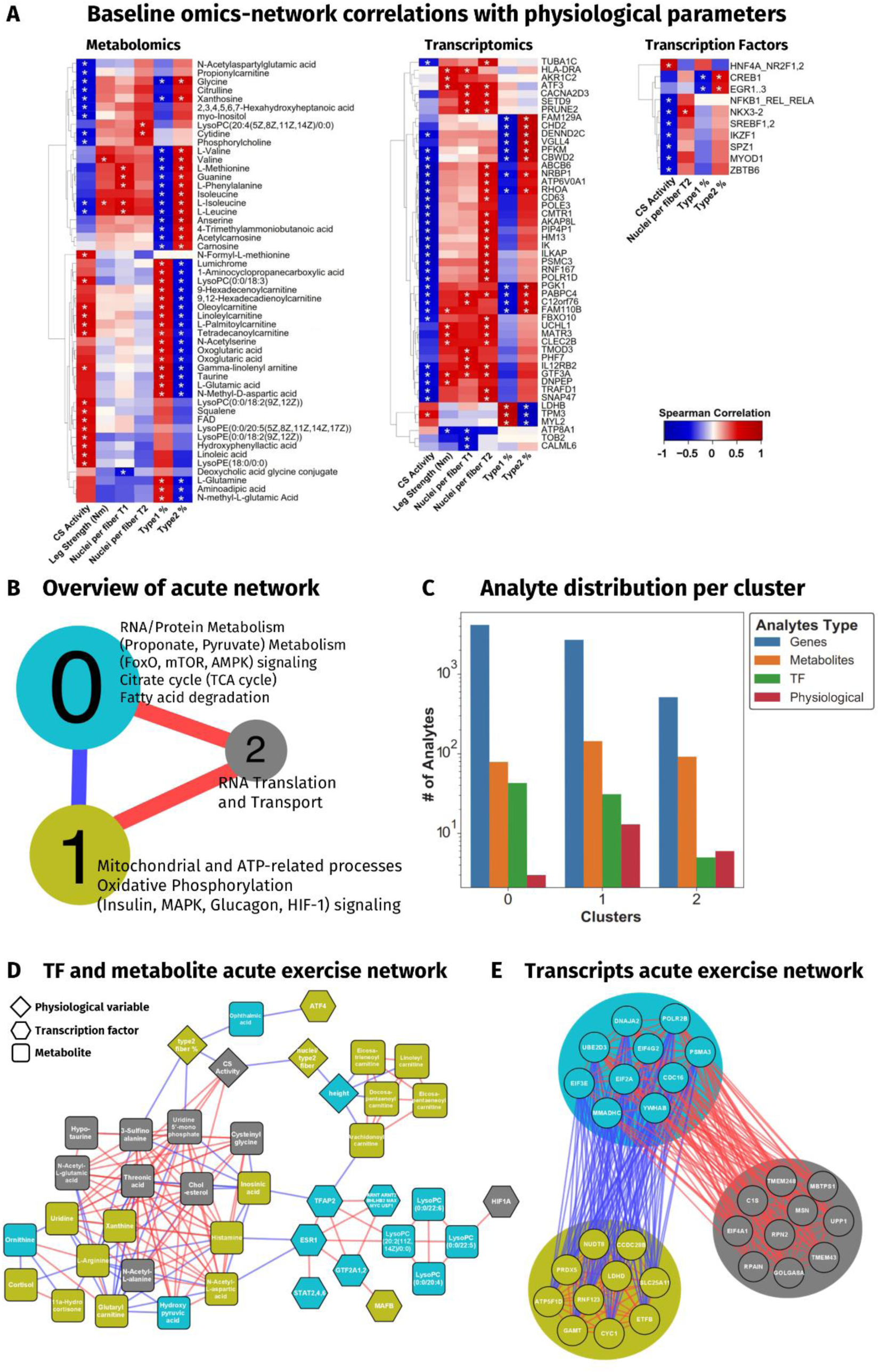
Omics-network overview and highlights. (A) Baseline -omics network showing the up to top 10 correlating genes for each physiological variable and all correlating transcription factor motifs and metabolites. (B) Overview over the acute -omics network with functional annotations of the three main clusters. (C) Distribution of analytes within the individual clusters from C. (D) Network of the top 10% of metabolites, transcription factor motifs and physiological variables from each cluster. Edge colors represent direction and strength of connections. Red = positive, blue = negative. (E) Top 10 genes from each cluster, edges follow the same logic as in D. (fill colors in D and E represent cluster identity as in B; *: FDR<0.01 for Spearman correlation)

**Figure 6:**
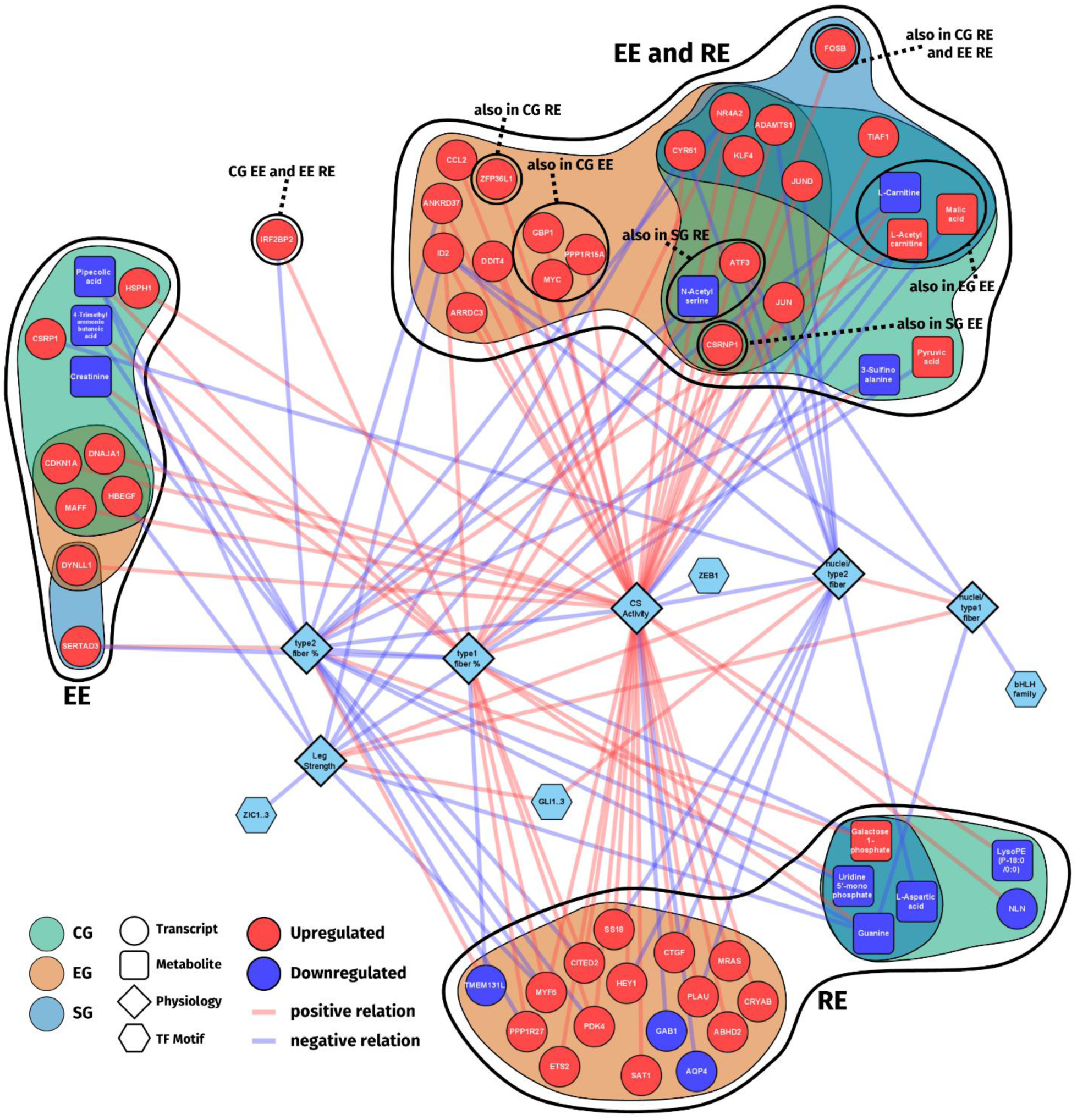
Multi-omics network of acute exercise. Multi-omics network of acute exercise showing genes, metabolites and transcription factor motifs significantly changed with acute exercise and positive or negative connection to selected physiological variables grouped by form of acute exercise and subject group.

#### Omics-network at baseline

To analyze the network at baseline we selected physiological, biochemical and histological parameters such as leg strength, CS activity and fiber type composition and analyzed their inter-network connections. Results show a high number of metabolites correlating with the proportion of type 1 (T1%) and 2 fibers (T2%), particularly in the amino acid group, positively correlating with T2%, and fatty acid group of metabolites, correlating with T1% (Figure 5A). CS activity was positively correlated primarily with PC- and PE-compounds and four carnitine derivates, and negatively correlated with amino acids, nucleosides and propionylcarnitine. Such an increased presence of endogenous muscle fatty acids^72^, associated with an increase in beta-oxidative capacity has previously been demonstrated^73, 74^, representing a shift to a more fatty-acid based energy metabolism. Leg strength showed substantially less correlation to the metabolome at rest, however with the exception BCAAs valine and L-isoleucine. BCAAs are capable of activating mTORC1 and to stimulate muscle protein synthesis^75^, a key process for muscle hypertrophy, but are also relevant to muscle regeneration from acute exercise^76–78^. In the transcriptome, results showed a large negative correlation between gene expression and parameters associated with EG (CS activity and T1%), while gene expression was largely positively correlated with parameters associated with SG (leg strength, myonuclear number and T2%) (Figure 5A). This could be explained by the increased at-rest potential capacity of RNA synthesis in SG caused by an increased proportion of myonuclei per muscle fiber (Table 1). One notable exception to this trend of negative correlation in EG was TPM3, which was positively correlated with a greater T1% and previously reported as involved in the calcium-dependent regulation of slow-twitch skeletal muscle fiber, significantly contributing to their contractile force^79^. For transcription factor motifs we observed the same negative trend of most notably the CREB1 motif correlation with T1%, but an increased motif activity with T2% (Figure 5A).

#### Omics-network of the acute exercise response

To better define the structure of the acute network we created, we performed community detection analysis using Leiden clustering algorithm and identified three main clusters after maximizing the modularity score of all types of analytes (Figure 5B). This approach provides a better understanding of the general processes taking place in our created network. Results showed the biggest cluster (cluster 0) to be associated with RNA and protein metabolism, FoxO, mTOR and AMPK signaling, the TCA cycle and fatty acid degradation. In animal models, FoxO signaling has previously been shown to improve glucose and lipid metabolic control via gluconeogenic enzymes^80–83^ and lipase activity^80^, consequently increasing metabolic control. Furthermore, it has been implicated in the process of muscle mass maintenance and in controlling the myogenic factors in humans, extensively reviewed by Gross et al^84^. Similarly, mTOR is a key mediator of muscle protein synthesis^85, 86^ and is stimulated by resistance exercise to a higher extent than by endurance exercise^87^. AMPK is a metabolic master switch, downregulating anabolic-, and upregulating catabolic processes, as well as inducing mitochondrial biogenesis^88, 89^. Also, it is influenced by being inhibited via acute exercise-reduced phosphocreatine^90^, but also long-term via transcription factors such as, GEF and MEF2 and the transcriptional coactivator PPARGC1A^91^. In combination with the other functions of cluster 0, these key regulatory axes can dictate metabolic status and trajectory in skeletal muscle. Cluster 0 was negatively correlated to cluster 1, which was associated with mitochondrial and ATP-related processes, oxidative phosphorylation and insulin-, MAPK-, glucagon- and HIF-1 signaling. Again, energy metabolism is central to this cluster, with anabolic, catabolic and amphibolic elements, such as MAPK which acts via targets such as PPARGC1A, ATF2 and CREB^92^. In addition to the above-mentioned systemic role of HIF-1A, locally it can indicate increased hypoxia as a consequence of increased oxygen-consuming ATP provision. Both these clusters were positively correlated to cluster 2, associated with RNA transport and translation into proteins. Due to the overrepresentation of transcriptomic analytes (Figure 5C), analytes with the highest degree of connection are shown separated into top 10% TFs and metabolites (Figure 5D), and top 10 transcripts (Figure 5E). Here, metabolites of cluster 2 showed a higher degree of centrality, supporting cluster 2’s central role in driving protein translation for pathways of both the other clusters. Furthermore, on individual analyte level, reiterating the most central pathways of cluster 1 in Figure 5B, six of the top 10 genes (SLC25A11, NUDT8, LDHD, ATP5F1D, CYC1, ETFB) of cluster 1 are mitochondrial protein-coding genes that are part of the electron transport chain or the TCA cycle.

To contextualize the acute exercise network with measures of performance, we selected EG- and SG-representative parameters such as CS activity, leg strength and T1% and T2%. Then we selected only analytes (transcripts, metabolites, TFs) that were significantly correlated with these parameters, filtered the resulting network with only analytes that were significantly changed with acute exercise and grouped them by acute exercise and subject group (Figure 6). This resulted in the visualization of genes that are both, significantly affected by training background and acute exercise, reinforcing their high relevance for exercise training. Most analytes were connected to CS activity or T2%, the two physiological parameters with the highest degree of centrality. EG showed to be the most affected group, specifically in the type of exercise they were not used to (RE). In a functional analysis using gene ontology in EG, exclusively RE stimulated genes significantly associated with anatomical structure formation involved in morphogenesis and the response to hormones. Most notable were previously mentioned MYF6, CITED2, an inhibitor of HIF1A signaling and modulator of PPARGC1A^93–95^, and PDK4, a mitochondrial protein that redirects the metabolic flux from glycolysis to fat metabolism^96^. This high number of RE responsive genes in EG compared to SG suggest a higher potential transcriptomic stimulation potential in response to the form of exercise they are not used to. Further, the stimulated transcripts of EG in response to RE are primarily associated with CS activity rather than RE-related performance parameters such as T2%. This suggests a fundamentally different molecular response when performing RE in endurance trained compared to strength trained individuals. However, 5 core genes for both EE and RE in all three groups were revealed in this approach. These include three transcription factor related genes (JUND, KLF4, NR4A2) and two genes that are involved in growth factor signaling, CCN2 and AMATS1, the latter which has been shown to interact with VEGFA^97^, suggesting a common, exercise-independent growth and transcription factor signaling program.

## DISCUSSION

Multi-omics network integration analysis of the molecular response to acute endurance and resistance exercise revealed distinct differences between high-level athletes, particularly endurance athletes, and control subjects, but also between the two exercise types. We show that life-long, high-level training results in a divergent, exercise-specific specialization of molecular mechanisms, resulting in individually rearranged acute molecular exercise responses.

### Training-background specific adaptations of energy metabolism

A key adaptation to exercise training lies in the way ATP is produced, which in EG is reflected in the enrichment of mitochondrial and energy metabolic pathways at baseline compared to both the other groups, a consequence of their increased CS activity and mitochondrial volume. On metabolic level, carnitines, chaperone molecules of beta-oxidation, and fatty acids of different lengths bound to these carnitines were enriched in EG, while SG-enriched PCs and PEs contribute to membrane fluidity and dynamics of, amongst others, GLUT4, which improves glucose uptake potential. Such metabolic adaptations reflect an improvement of the commonly used energy metabolic systems – with beta-oxidation improvements in EG and glucose metabolism improvements in SG. Unexpectedly, analysis of the acute exercise response revealed a steep downregulation of both energy metabolic pathways and transcriptomics-based metabolic flux in EG only. Previous studies have show EG adaptation to largely lie in energy metabolic optimization, providing a high flux of ATP over periods of extended exercise^98^. Thus, a sharp immediate decrease in energy metabolic pathways upon cessation of acute exercise suggests a tight regulatory control as a consequence of this adaptational goal. This notion is additionally supported by EG’s and SG’s metabolic response to EE and RE, respectively. Specifically, emphasizing carnitine metabolism in endurance athletes performing EE and AA metabolism in resistance-trained athletes performing RE suggests a metabolic specialization to the familiar form of exercise. In EG, carnitine-derivates are largely upregulated following EE, and largely downregulated following RE, a distinction not found in CG and SG. In SG, AAs are primarily downregulated performing RE, but not following EE, while downregulation also occurs in EG doing EE. These adaptations allow for physical performance beyond the levels of untrained individuals. One example of the ad-hoc upregulation of energy flux common across all groups is the metabolite sorbitol, aiding glucose uptake into the cell, which we, for the first time in humans, report to be upregulated in response to EE and RE. In our integrated network analysis, further evidence highlights the core role of energy metabolism. Energy metabolism is a central player in both forms of acute exercise – where the citrate cycle, FA and protein metabolism, and insulin and glucagon signaling are prominently represented in cluster annotations (Figure 5B). Energy metabolism-related group differences can further be observed in the at-rest network, with carnitine metabolites correlating with high aerobic capacity, a proxy for endurance performance, and amino acid metabolites correlating with strength parameters. Particularly, one of the genes exclusively regulated in EG in response to RE was PDK4, a gene responsible for orchestrating metabolic flux, further reinforcing the notion of the EG specialization towards optimization of energy production. Together, this suggests exercise-type metabolic adaptation and different degrees of energy metabolic control, stemming from accumulation of repeated bouts of acute specific exercise, adding the facet of acute exercise to our understanding of endurance training adaptation and underlining the substantial uniqueness of EG.

### Adaptational potential is retained in highly specialized athletes

Several findings in this current study but also previously published results support the notion of a retained high potential for resistance exercise adaptation in EG compared to SG: Previous publications demonstrated that exercise-naïve individuals exhibit a prolonged elevation of muscle protein synthesis following a bout of acute resistance exercise^99^, and that previous resistance exercise experience increasingly protects against muscle damage from subsequent bouts^71^, laying the foundation for the idea of a retainment of adaptational potential even in highly specialized, though RE-naïve EG. In our results of the created acute network we showed high numbers of genes exclusively regulated in EG in response to RE which are highly relevant for RE-specific adaptation processes. Apart from the above mentioned PDK4 it also included MYF6 and CITED2, regulating hypertrophy and angiogenesis (Figure 6). Additionally, all groups significantly increased HIF1A and MYFfamily TF activity in response to EE and RE, respectively. This is noteworthy because both of their gene targets are early activated in the regulatory cascade of exercise-specific adaptation^52, 53^. Beyond this, following an acute bout of exercise they are not adapted to, both athlete groups retain a higher number of DEGs. Additionally, regulation of EE and RE follows an exercise type-specific pattern, with an increase in HIF1A and MYFfamily TF motif activity respectively, despite the above-mentioned group-specific specializations. This indicates that the form of acute exercise rather than the training history is the primary determinant of the individual exercise session response.

### Life-long training induces molecular specialization by transcriptional narrowing and amplification

Gene clusters respond similarly to the form of acute exercise regardless of subject group, however to a different extent, with a substantially higher effect amplitude in the familiar form of exercise (Figure 2C). However, at the 3h timepoint, we found a *lower* number of DEGs in athletes performing their accustomed form of exercise compared to their exercise-naïve peers (Figure 1C). While it might be tempting to accredit this to a simple blunting of gene expression effect, together with above mentioned increase in the response amplitude, this can also be explained by an optimized response in the respective experienced athletes. Following acute exercise, DEGs increase from post to the 3h timepoint and are increasingly dominated by regeneration-related pathways. A commonly observed effect of long-term training is an optimized regeneration from acute exercise^100^, which might cause such a reduced number of DEGs. However, we showed the gene clusters in response to EE and RE to be strongly associated with more adaptational pathways such as transcriptional regulation and translation, and muscle filament-related processes. Additionally, RE only was associated with myoblast differentiation, growth, and morphogenesis. Taken together, these observations suggest the difference between subject groups is based on the degree of gene activation rather than number of DEGs. This is in line with a previous publication that indicated muscle memory effects to be mediated by modification of the translational capacity, and thus potential amplitude of protein expression in the context of acute exercise^101^. Additional observations support the idea of an exercise-specific trajectory of development from untrained to trained state via degree of molecular and metabolic exercise-specific optimization: On metabolomic level in EG, lipid-associated metabolites were reduced in response to RE but not EE (Figure 3A). In SG however, RE largely decreased amino acid-instead of lipid-associated metabolites, indicating a lower usage of lipids as a source of energy in SG.

### Performance diversity can be explained by small numbers of genes

By correlating gene expression and physiological performance parameters, we showed, that only 4 genes can explain 68% of the leg strength variation and 5 genes 42% of V̇O2peak. Interestingly, GSEA analysis associated the RE-correlating genes with energy metabolism and transcription and translation processes. Despite being less studied, RE requires and results in substantial energy metabolic perturbations^102^. Together, this might underline the importance of further research into the resistance exercise-related energy metabolism. On the other hand, the broad influence of V̇O2peak results in a higher *number* of significantly correlating genes, mainly associated with transcription processes and their regulation. Leg strength correlates stronger with individual genes as it is a closer representation of *local* performance and thus more adequately representing *local* gene expression, in particular energy metabolic pathways important for and perturbed by acute exercise.

### In-silico methods add to the understanding of complex data

Metabolic flux related to the citrate cycle and beta-oxidation showed to be distinctively down-regulated in EG only following EE as analyzed by genome-scale metabolic modelling (GEM), indicating EGs particularly tight metabolic control over energy metabolic pathways as discussed above. Additionally, we identified HES1, CLIC4 and TRIM63 as key targets of HIF1A and MYFfamily TF motifs following EE and RE respectively, demonstrating a robust form of exercise-specific gene regulatory program independent of exercise background. These findings demonstrate the ability of both these in-silico methods to enhance the perspective on transcriptomic data by adding database-originating layers of information.

## Conclusions

Multi-omics and large data approaches are increasingly used for precision health and physical activity research. Here, we employed such an approach to compute differences between endurance and strength athletes and untrained individuals, showing major differences in energy metabolic and amino acid-related pathways following acute exercise. Highly endurance trained athletes specialize in lipid-related energy processes while resistance trained athletes change the way they utilize AAs, a diverging development from the untrained state visible from omics and network perspectives. We further enhanced our analysis using GEM and MARA analysis, showing a striking down-regulation of metabolic flux as a consequence of EGs training-induced metabolic adaptation and identifying the HIF1A and MYFfamily TF motif as exclusive to EE and RE respectively, independent of exercise background.

Together, these findings put forward the concept of a numerically smaller but more intense stimulation of genes – a transcriptional specialization - with life-long high-level training, a metabolic specialization focused on the needs of the type of exercise training performed, and a certain degree of transcription factor motif specificity to the form of acute exercise, driven by particular sets of exercise-responsive genes. Finally, our transcriptomic, metabolomic and regulomic data can be a useful resource for further investigations into the molecular biology of exercise.

## ACKNOWLEDGEMENTS

The authors would like to thank the Division of Clinical Physiology at Karolinska Universitetssjukhuset Huddinge for providing the clinical research environment to conduct this study. The authors acknowledge support from the National Genomics Infrastructure Stockholm funded by Science for Life Laboratory, the Knut and Alice Wallenberg Foundation and the Swedish Research Council. The Swedish Metabolomics Centre, Umeå, Sweden (www.swedishmetabolomicscentre.se) is acknowledged for metabolic profiling by GC/LC-MS. Imaging was performed in the Biomedicum Imaging Core with support from Karolinska Institutet. We would also like to thank the Swedish Research Council (VR, grant number 2018-02932), the Swedish Research Council for Sport Science (CIF) and the research subjects volunteering to be part of this study.

## METHODS

### KEY RESOURCES TABLE

| REAGENT or RESOURCE | SOURCE | IDENTIFIER |
| --- | --- | --- |
| **Antibodies** |  |  |
| Anti-Laminin antibody produced in rabbit | Sigma-Aldrich | Cat# L9393, RRID: AB_477163 |
| Anti-Myosin heavy chain type 1 | DSHB | Cat# BA-F8,RRID: AB_10572253 |
| Anti-Myosin heavy chain type 2 | DSHB | Cat# SC-71, RRID: AB_2147165 |
| Goat Anti-Rabbit IgG (H+L) Antibody, Alexa Fluor 488 Conjugated | Molecular Probes | Cat# A-11008, RRID:AB_143165 |
| **Chemicals, Peptides, and Recombinant Proteins** |  |  |
| <u>Reference and tuning standards:</u> |  |  |
| LC/MS reference mass standard kit | Agilent Technologies | Part# G1969-85003 |
| HP-0921 (Hexakis(<sup>1</sup>H, <sup>1</sup>H, <sup>3</sup>H-tetrafluoropropoxy)phosphazine) | Agilent Technologies | Part# G1969-85001 |
| ESI-TOF, ESI-L Low Concentration Tuning Mix | Agilent Technologies | Part# G1969-85000 |
| <u>LC-MS internal standards:</u> <sup>13</sup>C<sub>9</sub>-Phenylalanine, <sup>13</sup>C<sub>3</sub>-Caffeine, D<sub>4</sub>-Cholic acid, D<sub>8</sub>-Arachidonic Acid, <sup>13</sup>C<sub>9</sub>-Caffeic Acid | Sigma | Product# 795844, 902322, 614149, 735000, 706337 |
| <u>GC-MS internal standards:</u> L-proline-<sup>13</sup>C<sub>5</sub>, alpha-ketoglutarate-<sup>13</sup>C<sub>4</sub>, myristic acid-<sup>13</sup>C<sub>3</sub>, cholesterol-D<sub>7</sub> | Cambridge Isotope Laboratories | Item# CLM-2260-H, CLM-4442, CLM-3665, DLM-3057 |
| <u>GC-MS internal standards:</u> Succinic acid-D<sub>4</sub>, salicylic acid-D<sub>6</sub>, L-glutamic acid-<sup>13</sup>C<sub>5</sub>, <sup>15</sup>N, putrescine-D<sub>4</sub>, hexadecanoic acid-<sup>13</sup>C<sub>4</sub>, D-glucose-<sup>13</sup>C<sub>6</sub>, D-sucrose-<sup>13</sup>C<sub>12</sub> | Sigma | Product# 293075, 616796, 607851, 51799, 489611, 389374, 605417 |
| **Deposited Data** |  |  |
| RNA sequencing data | This manuscript | EGA: EGAS00001006139 |
| **Software and Algorithms** |  |  |
| Zeiss Zen Pro v2.0 | Zeiss International | https://www.zeiss.com/microscopy/int/products/microscope-software/zen.html |
| ImageJ | NIH, https://imagej.net | RRID:SCR_003070 |
| RStudio 1.3.1056 | https://rstudio.com | RRID:SCR_000432 |
| R 3.6.0 | https://r-project.org | RRID:SCR_001905 |
| **Other** |  |  |
| Mounting medium | Vectashield | Cat# H-1000 |

### RESOURCE AVAILABILITY

#### Lead contact

Further information and request for resources and reagent should be directed to and will be fulfilled by the lead contact, Stefan Markus Reitzner.

#### Materials Availability

This study did not generate new unique reagents.

#### Data and Code Availability

The RNA-sequencing data has been deposited at the European Genome-phenome Archive (EGA) which is hosted at the EBI and the CRG, under accession number EGA: EGAS00001006139. Metabolomics data is available as supplement to this paper.

### EXPERIMENTAL MODEL AND SUBJECT DETAILS

#### Ethical Approval

Before the interventions, all subjects were informed about the study outline, familiarized with the experimental procedures and potential complications and written and verbal consent was obtained. The study was approved by the Stockholm regional ethics board (Dnr: 2016/590-31) under observance of the declaration of Helsinki.

#### Participant Recruitment and Inclusion

Twenty-four healthy, non-smoking men aged 33-52 were recruited after filling in a questionnaire to assess their training history and health status and a selection process including peak oxygen uptake test (V̇O_2_peak test) and maximum knee torque test measured by unilateral isokinetic Biodex test (Biodex System 4, Biodex Medical Systems, Shirley (NY), USA) to fit into one of three groups: resistance trained, endurance trained and control. V̇O_2_peak test was performed on a stationary bike and initial resistance individually set depending on expected subject performance. After 5 minutes of warm-up cycling, resistance was increased incrementally between 16.6 and 26.6 W⋅min^-1^ until exhaustion. All subjects reaching a respiratory exchange ratio (RER) above 1.05. Inclusion into the EG required a V̇O2peak above the 90^th^ percentile of the subject’s age group^103^. Additionally, knee extension 1 repetition maximum (1RM) was measured and controlled using the Borg RPE scale^104^. Inclusion into the SG required a peak torque of at least two standard deviations above the mean of the control group (Table 1). The resistance and endurance trained athletes were required to have either a resistance- or an endurance-based training history at high level of at least 15 years respectively. These groups were clearly separated by both, V̇O2peak and maximum knee torque (Table 1). Thresholds for V̇O2peak and peak torque were selected to be identical to our previously published cross-sectional study to ensure maximal comparability^10^.

### METHOD DETAILS

#### Study Design

All subjects completed both a single bout of EE and RE with at least one month of wash-out in between (**Error! Reference source not found.**A). Food intake was controlled for by a standardized breakfast and the instruction to eat the same dinner on the day before each of the intervention days. After a standardized breakfast 3 hours prior to the first biopsy timepoint, subjects arrived between 8:00 and 9:00 in the morning at the Division for Clinical Physiology research unit at the Karolinska Hospital in Huddinge, Sweden. Subjects were then randomized to either an acute endurance or acute resistance exercise session. After a resting skeletal muscle biopsy was taken using the Bergström needle technique with suction from *M. vastus lateralis*, subjects performed 30 minutes of acute exercise. Subjects were pushed to their maximum performance in each session. For acute endurance exercise, subjects were required to cycle at 75% of their individual W_max_ for 30 minutes. For acute resistance exercise, subjects were required to complete 9 sets of 8 repetitions of knee extension (set length 40 seconds; set break, 150 seconds) exercise at 80% of their individual one repetition maximum. Following the conclusion of the acute exercise, three additional skeletal muscle biopsies were taken from *M. vastus lateralis* immediately following acute exercise and 1 hour and 3 hours after the end of the acute exercise. Between 4 and 8 weeks after the first session, all subjects returned to perform the form of exercise they have not been randomized for the first occasion and skeletal muscle biopsies were again collected at a total of four timepoints.

#### Skeletal muscle tissue biopsies

Skeletal muscle biopsies were taken alternating from left and right M. vastus lateralis before and directly after and at 1 hours and 3 hours after one single bout of exercise using the Bergström needle technique^105^. To prevent possible dominant/non-dominant leg effects, the first muscle biopsy was taken from either the dominant or the non-dominant leg by randomization. The skeletal muscle biopsies were then snap-frozen using 2-methylbutane as a secondary coolant.

#### Baseline tissue characterization

##### Enzyme extraction and enzymatic assays

Enzymes were extracted using a phosphate-based buffer (50 mM KH2PO4, 1 mM EDTA, 0.05 % Triton X-100) and quantified by Bradford assay (Bio-Rad #5000006, Bio-Rad, Hercules, CA, USA). Citrate synthase activity was measured spectrophotometrically as previously described^106^.

##### Immunohistochemical fiber typing

Skeletal muscle samples were cut on a cryostat (CryoStar NX70, Thermo Scientific, Waltham, MA, USA) at 7 µm and immunohistochemically stained for Type I (DSHB, Cat# BA-F8, RRID:AB_10572253) and Type II (DSHB, Cat# SC-71, RRID:AB_2147165) fibers and Laminin (Sigma-Aldrich, Cat# L9393, RRID:AB_477163). Compatible fluorescent antibodies were used for detection with the Olympus IX-73 wide-field fluorescence microscope (Olympus Corp., Shinjuku, Tokyo, Japan). Analysis of immunohistochemically stained cross-sectional cuts was performed blinded using ImageJ.

#### RNA sequencing

RNA extraction from skeletal muscle tissue was performed using the phenol based TRIzol method (Invitrogen #15596018, Thermo Fisher Scientific, Waltham, MA, USA), quantified and quality checked using the 2100 Bioanalyzer System (Agilent Technologies, Santa Clara, CA, USA). Libraries were prepared by poly-A selection (TruSeq mRNA, Illumina, San Diego, CA, USA) and multiplexed at the National Genomics Infrastructure Sweden. Clustering was done by ’cBot’ and samples were sequenced on NovaSeq6000 (NovaSeq Control Software 1.6.0/RTA v3.4.4) with two lanes of a 2×151 setup ’NovaSeqXp’ workflow in ’S4’ mode flow cell. The Bcl to FastQ conversion was performed using bcl2fastq_v2.20.0.422 from the CASAVA software suite. The quality scale used is Sanger / phred33 / Illumina 1.8+. QC and processing were performed using the nfcore/rnaseq analysis pipeline publicly available at github (https://github.com/nf-core/rnaseq).

#### Semi-targeted Metabolomics by Gas- and Liquid Chromatography (GC/LC)-MS

Metabolic profiling by GC-MS and LC-MS was performed at the Swedish Metabolomics Center in Umeå, Sweden.

##### Sample Preparation

Sample preparation of 20 mg of skeletal muscle tissue was performed according to A et al.^107^. In detail, 1000 µL of extraction buffer (80/20 v/v methanol : water) including internal standards for both GC-MS and LC-MS were added to the tubes together with 2 tungsten beads. The sample was shaken at 30 Hz for 3 minutes in a mixer mill. The sample was centrifuged at +4 °C, 14 000 rpm, for 10 minutes. The supernatant, 200 µL for LCMS analysis and 50 µL to GCMS analysis, was transferred to micro vials and evaporated to dryness in a speed-vac concentrator. Solvents were evaporated and the samples were stored at −80 °C until analysis. A small aliquot of the remaining supernatants was pooled and used to create quality control (QC) samples. MSMS analysis (LCMS) was run on the QC samples for identification purposes. The samples were analyzed in batches according to a randomized run order on both GC-MS and LC-MS.

##### GC-MS Analysis

Derivatization and GC-MS analysis were performed as described previously^108^. 0.5 μL of the derivatized sample was injected in splitless mode by a L-PAL3 autosampler (CTC Analytics AG, Switzerland) into an Agilent 7890B gas chromatograph equipped with a 10 m x 0.18 mm fused silica capillary column with a chemically bonded 0.18 μm Rxi-5 Sil MS stationary phase (Restek Corporation, U.S.). The injector temperature was 270 °C, the purge flow rate was 20 mL⋅min^-1^ and the purge was turned on after 60 seconds. The gas flow rate through the column was 1 mL⋅min^-1^, the column temperature was held at 70 °C for 2 minutes, then increased by 40 °C min^-1^ to 320 °C and held there for 2 minutes. The column effluent was introduced into the ion source of a Pegasus BT time-of-flight mass spectrometer, GC/TOFMS (Leco Corp., St Joseph, MI, USA). The transfer line and the ion source temperatures were 250 °C and 200 °C, respectively. Ions were generated by a 70 eV electron beam at an ionization current of 2.0 mA, and 30 spectra s-1 were recorded in the mass range m/z 50 - 800. The acceleration voltage was turned on after a solvent delay of 150 seconds. The detector voltage was 1800-2300 V.

##### LC-MS Analysis

Before LC-MS analysis the sample was re-suspended in 10 + 10 µL methanol and water. Each batch of samples was first analyzed in positive mode. After all samples within a batch had been analyzed, the instrument was switched to negative mode and a second injection of each sample was performed. The chromatographic separation was performed on an Agilent 1290 Infinity UHPLC-system (Agilent Technologies, Waldbronn, Germany). 2 μL of each sample were injected onto an Acquity UPLC HSS T3, 2.1 x 50 mm, 1.8 μm C18 column in combination with a 2.1 mm x 5 mm, 1.8 μm VanGuard precolumn (Waters Corporation, Milford, MA, USA) held at 40 °C. The gradient elution buffers were A (H2O, 0.1 % formic acid) and B (75/25 acetonitrile:2-propanol, 0.1 % formic acid), and the flowrate was 0.5 mL⋅min^-1^. The compounds were eluted with a linear gradient consisting of 0.1 - 10 % B over 2 minutes, B was increased to 99 % over 5 minutes and held at 99 % for 2 minutes; B was decreased to 0.1 % for 0.3 minutes and the flow-rate was increased to 0.8 mL⋅min^-1^ for 0.5 minutes; these conditions were held for 0.9 minutes, after which the flow-rate was reduced to 0.5 mL⋅min^-1^ for 0.1 minutes before the next injection. The compounds were detected with an Agilent 6546 Q-TOF mass spectrometer equipped with a jet stream electrospray ion source operating in positive or negative ion mode. The settings were kept identical between the modes, with exception of the capillary voltage. A reference interface was connected for accurate mass measurements; the reference ions purine (4 μM) and HP-0921 (Hexakis(1H, 1H, 3H-tetrafluoropropoxy)phosphazine) (1 μM) were infused directly into the MS at a flow rate of 0.05 mL⋅min^-1^ for internal calibration, and the monitored ions were purine m/z 121.05 and m/z 119.03632; HP-0921 m/z 922.0098 and m/z 966.000725 for positive and negative mode respectively. The gas temperature was set to 150°C, the drying gas flow to 8 L⋅min^-1^ and the nebulizer pressure 35 psig. The sheath gas temp was set to 350°C and the sheath gas flow 11 L⋅min^-1^. The capillary voltage was set to 4000 V in positive ion mode, and to 4000 V in negative ion mode. The nozzle voltage was 300 V. The fragmentor voltage was 120 V, the skimmer 65 V and the OCT 1 RF Vpp 750 V. The collision energy was set to 0 V. The m/z range was 70 - 1700, and data was collected in centroid mode with an acquisition rate of 4 scans s-1 (1977 transients/spectrum).

##### Data Analysis and Evaluation

For the GC-MS data, all non-processed MS-files from the metabolic analysis were exported from the ChromaTOF software in NetCDF format to MATLAB R2016a (Mathworks, Natick, MA, USA), where all data pre-treatment procedures, such as base-line correction, chromatogram alignment, data compression and Multivariate Curve Resolution were performed. The extracted mass spectra were identified by comparisons of their retention index and mass spectra with libraries of retention time indices and mass spectra^109^. Mass spectra and retention index comparison was performed using NIST MS 2.0 software. Annotation of mass spectra was based on reverse and forward searches in the library. Masses and ratio between masses indicative of a derivatized metabolite were especially notified. If the mass spectrum according to SMC’s experience was with the highest probability indicative of a metabolite and the retention index between the sample and library for the suggested metabolite was ± 5 (usually less than 3) the deconvoluted “peak” was annotated as an identification of a metabolite.

For the LC-MS data, all data processing was performed using the Agilent Masshunter Profinder version B.08.00 (Agilent Technologies Inc., Santa Clara, CA, USA). The processing was performed both in a target and an untargeted fashion. For target processing, a pre-defined list of metabolites commonly found in plasma and serum were searched for using the Batch Targeted feature extraction in Masshunter Profinder. An-in-house LC-MS library built up by authentic standards run on the same system with the same chromatographic and mass-spec settings, were used for the targeted processing. The identification of the metabolites was based on MS, MSMS and retention time information. For the untargeted data, each tissue group was processed individually using Batch Recursive Feature Extraction algorithm within Masshunter Profinder.

### QUANTIFICATION AND STATISTICAL ANALYSIS

Statistical analyses were performed with R version 3.6.0. For baseline analyses, ANOVA and t-test were used. Gene expression was measured by differential gene expression determined by edgeR package analysis^110^. Analyses were also performed with subject groups or acute pooled to provide results for a background-independent general training effect. Repeated testing corrections were performed using the Benjamini-Hochberg method^111^. Using the data from differential gene expression analysis, gene set enrichment was determined by fGSEA based on log(FC)-sorting using the GO collections “cellular component”, “molecular function” and “biological process”. Gene and metabolite clustering was performed using k-means unsupervised clustering.

Area under der curve (AUC) for each gene was calculated using trapezoid integration and used for correlation analysis with baseline physiological performance markers VO_2_peak and peak torque using Pearson’s correlation coefficient. Modelling of top genes explaining VO2peak and peak torque was performed with linear modelling.

Genome-scale Metabolic Modelling (GEM) was performed using RNA sequencing results. Using GEMs, metabolites and metabolic pathways can be inferred by modelling metabolome information onto RNA sequencing results. Generic human model, HMR version 2.00 was used to generate gene-metabolite networks as previously described^36^.

Regulatory motif analysis was performed using a motif activity response analysis (MARA) algorithm^45^ and the FANTOM5 database of regulatory annotation^43, 44^. Of the 192 available motifs from the FANTOM5 annotation, 79 were left after running the MARA algorithm. These transcription factors motifs were assessed for their member genes that drive the temporal shape of the response to acute exercise. Only DEGs that were considered differentially regulated in the respective comparison in the differential gene expression analysis were considered for correlation to the motif using Pearson’s correlation coefficient. Verification of gene-gene interactions within regulatory motifs was performed using the tissue and cancer specific biological networks (TCSBN) database^112^. The multi-omics network was generated by using the same pipeline/method as described in iNetModels^112, 113^.

Alpha was set to 0.01, unless stated otherwise, error bars represent standard error of the mean.

## SUPPLEMENTARY FIGURES

**Figure S1:**
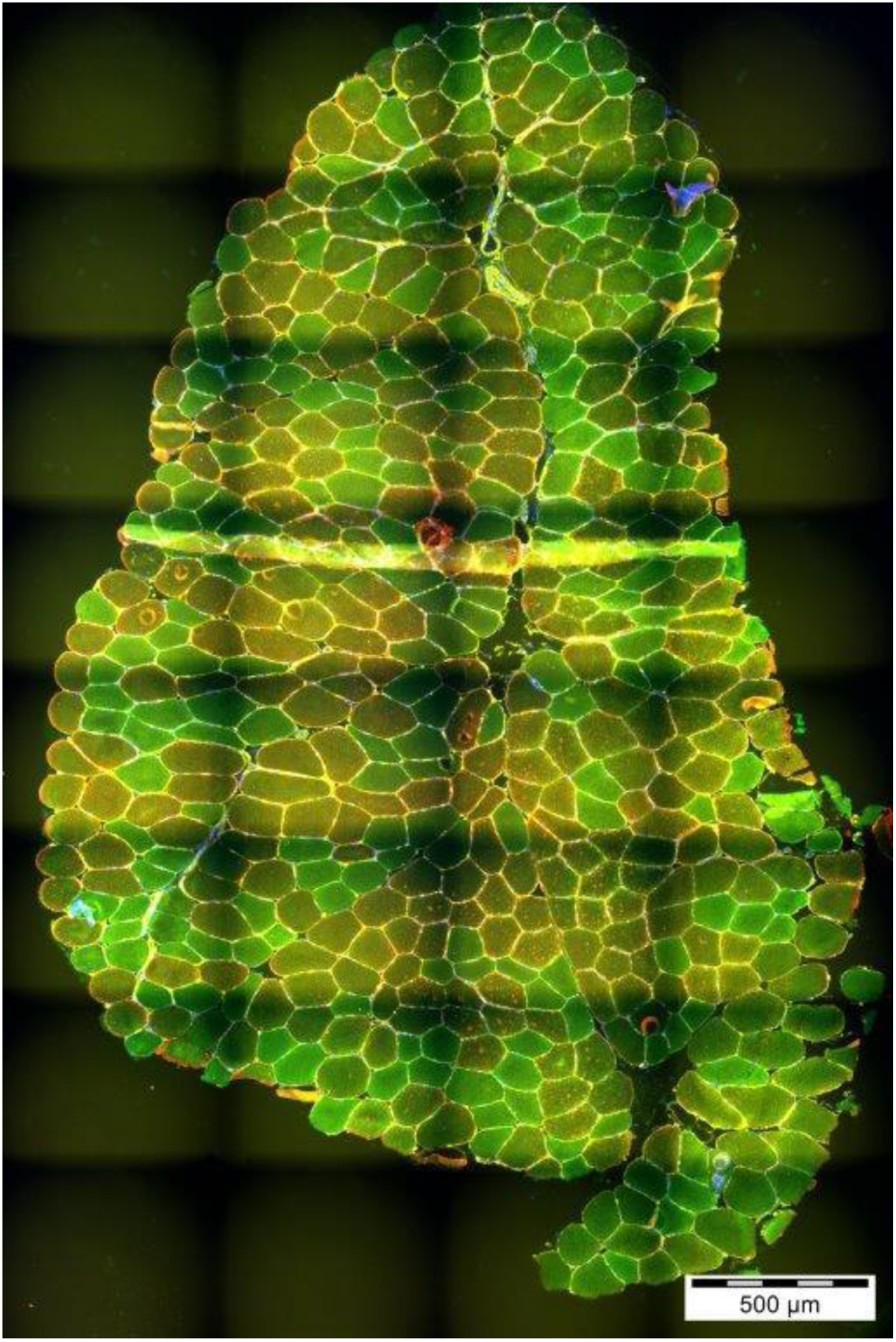
Histological analysis of skeletal muscle. Representative immunohistochemical staining of M. vastus lateralis. Staining was performed for Laminin (yellow), type I fibers (green) and type II fibers (red).

**Figure S2:**
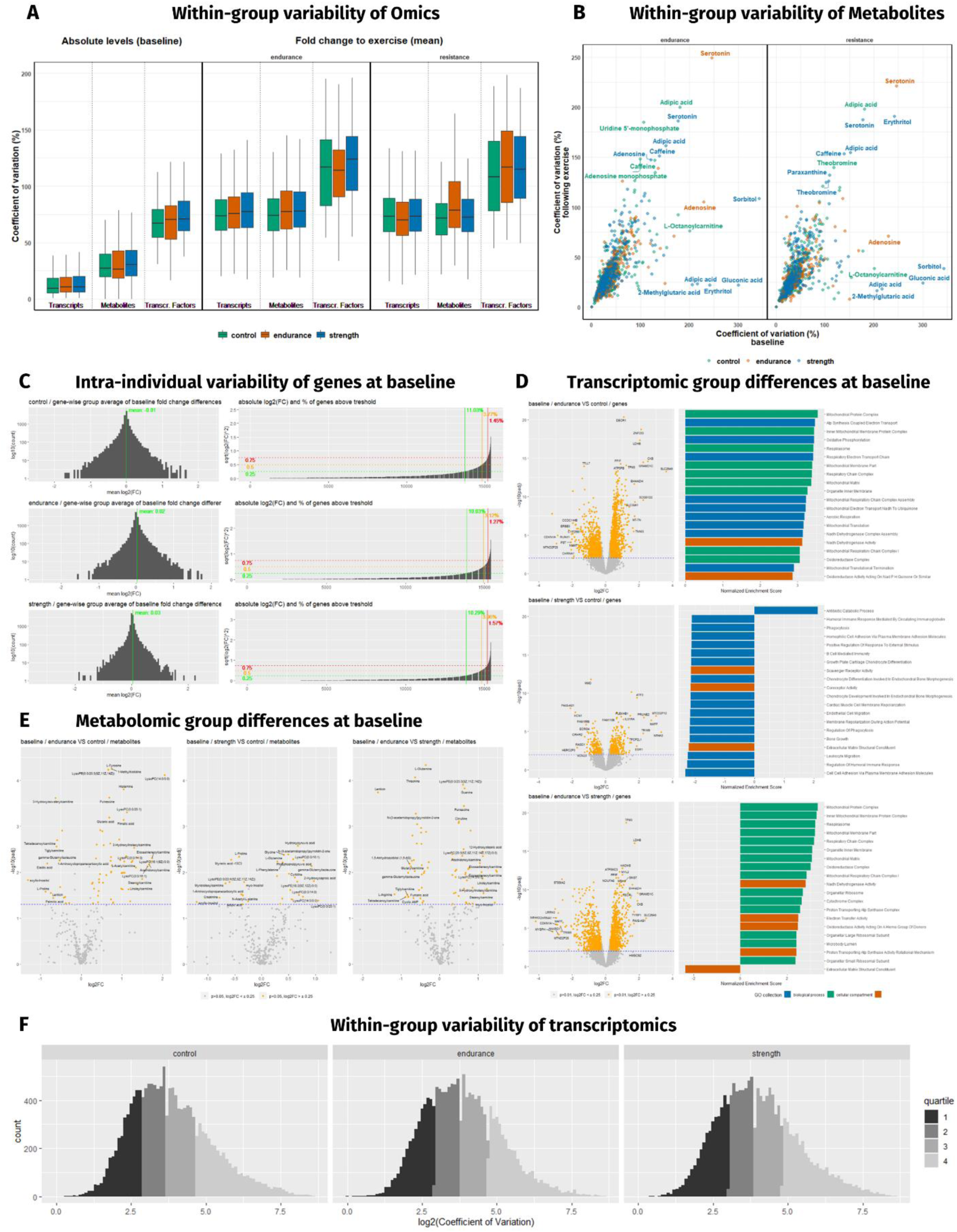
Intra-individual and -group differences and variation on transcriptomic and metabolomic level. (A) Within subject group variation of analytes (transcripts, directly measured metabolites and transcription factor motifs) at baseline and in response to acute endurance or resistance exercise (mean of time course). (B) Within subject group variation of metabolites at baseline (x-axis) and following acute exercise (mean of time course; y-axis) with the top metabolites in both dimensions annotated. (C) Intra-individual variability by gene-wise fold change analysis with mean and absolute fold change and proportion of genes above log fold change thresholds of 0.25, 0.5 and 0.75 separated by group. (D) Direct differential gene expression analysis of groups at baseline as volcano plot and the top 20 pathways of each gene set based on normalized enrichment score from gene set enrichment analysis. (E) Direct differential expression analysis of metabolites at baseline as volcano plot. (F) Density plot of the within-group coefficient of variation (CoV) for each gene separated by group. Quartiles of each set based on CoV are shown with different shades. CoV is presented as log2 transformation.

**Figure S3:**
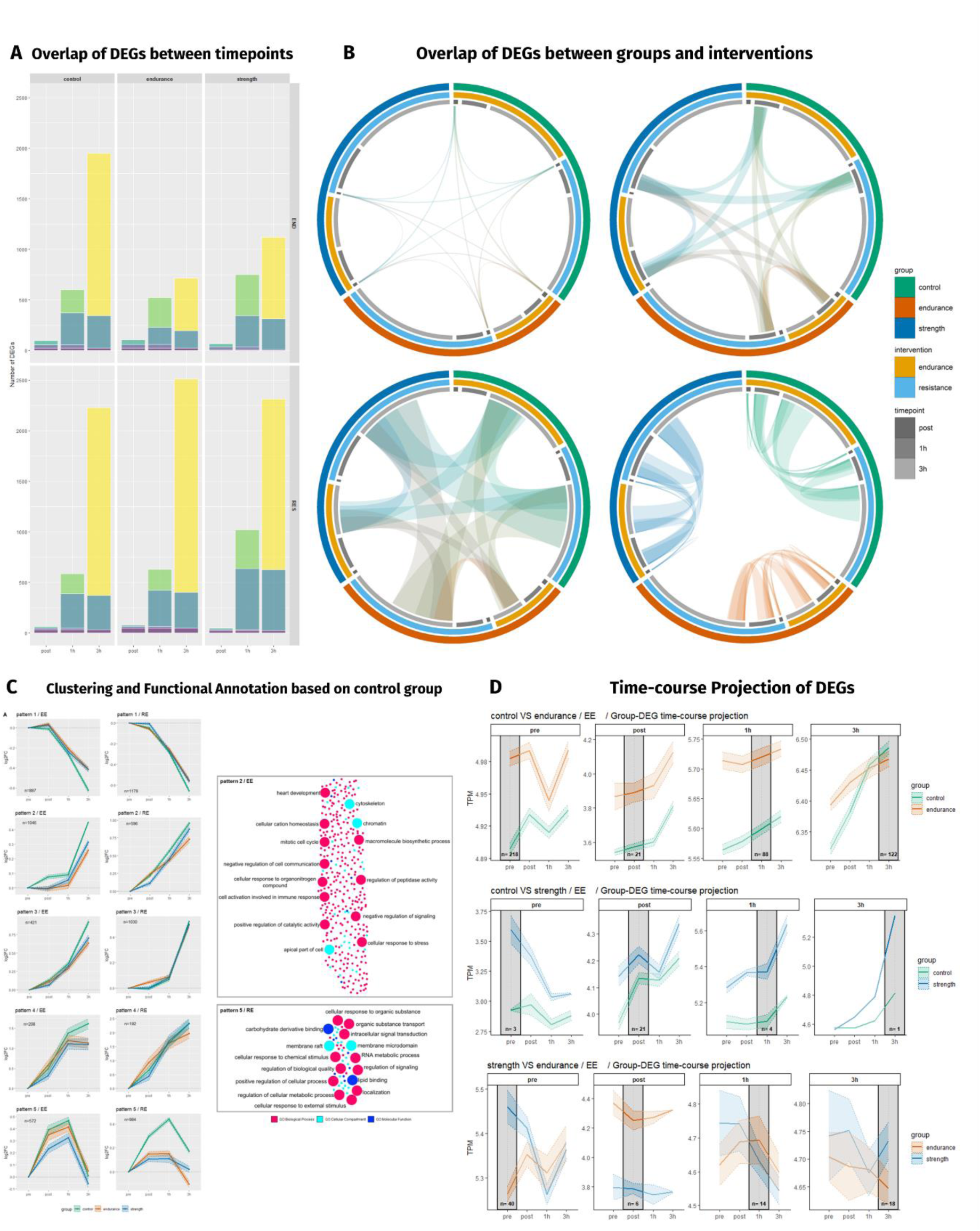
Analysis of DEG overlaps and functional annotation of clusters. (A) Overlaps of DEGs compared to pre-timepoint across the three post timepoints within control (CG), endurance (EG) and strength group (SG) in response to acute endurance (EE) and resistance exercise (RE). Colors represent sets of identical genes within each group-intervention pair. Genes common to all timepoints (core genes) ranged from 45 in EG performing RE (59% of all DEGs at post) to 6 in SG performing EE (9%). RE resulted in a higher proportion of core genes than EE across all groups. (B) Intra-group overlap of genes between groups and interventions at pre, post, 1h and 3h timepoints and of timepoints and acute interventions within groups. (C) Comparison of gene cluster trajectories based on unsupervised clustering of DEGs of the control group (CG) in response to both EE and RE. (D) Time-course projection (plotting of DEGs from one timepoint across the remaining timepoints) of genes significantly different between two groups at individual timepoints (solid line). Dotted lines show the confidence interval.

**Figure S4:**
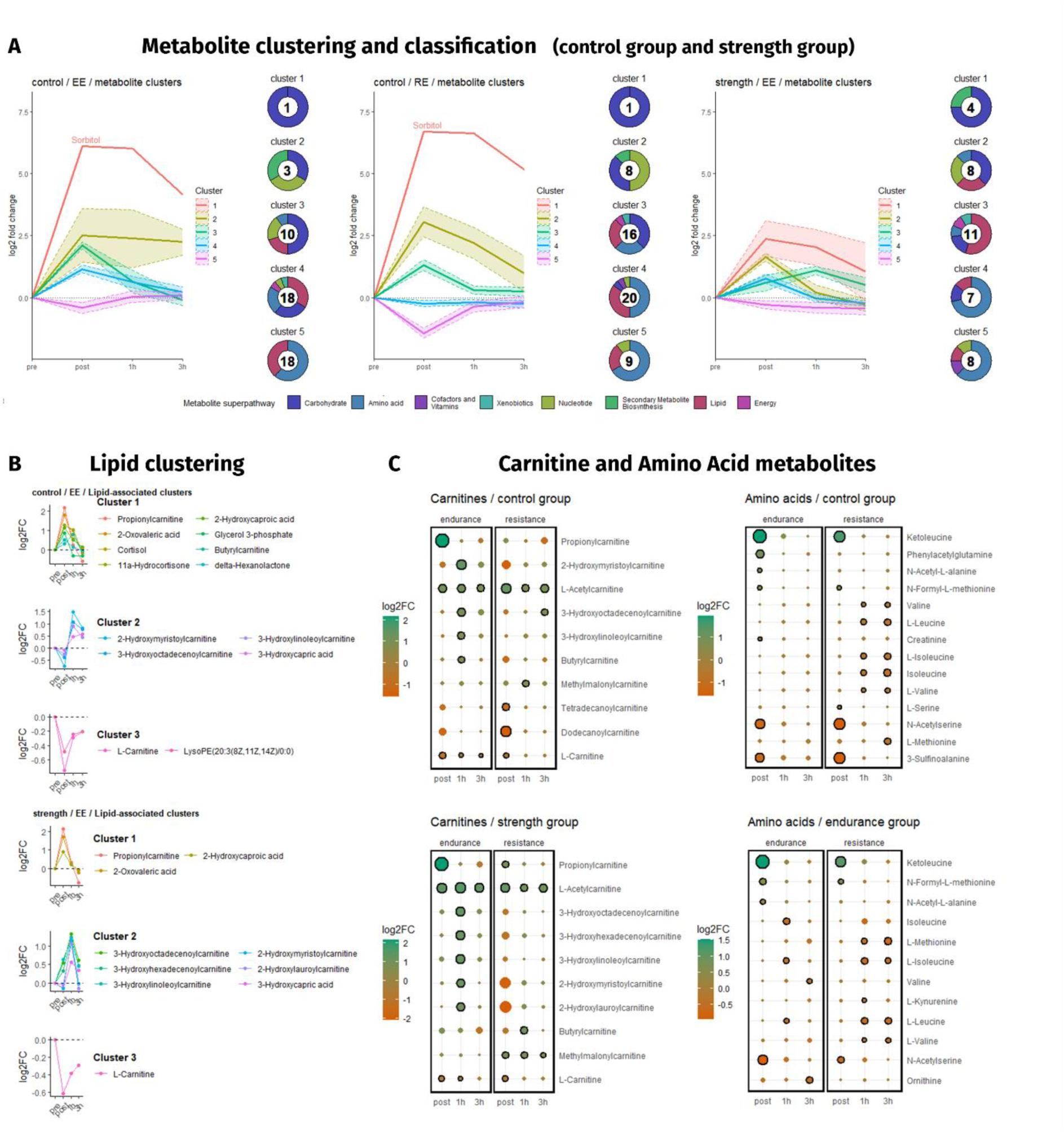
Clustering and classification of metabolites. (A) Clustering and classification of metabolites of control (CG) and strength group (SG) in response to acute endurance (EE) and resistance exercise (RE). Solid lines show cluster mean, dotted lines show the confidence interval. Numbers inside the donuts represent the number of metabolites in the cluster. (B) Isolated clustering of all metabolites associated with lipids or lipid metabolism in control (CG) and strength group (SG) in response to acute endurance exercise. (C) Analysis of carnitines in control (CG) and strength group (SG) and amino acids in control (CG) and endurance group (EG) in response to both forms of acute exercise. Color and size of dots represent effect size and direction. Dots with solid black circles are statistically significant timepoints compared to pre timepoint.

**Figure S5:**
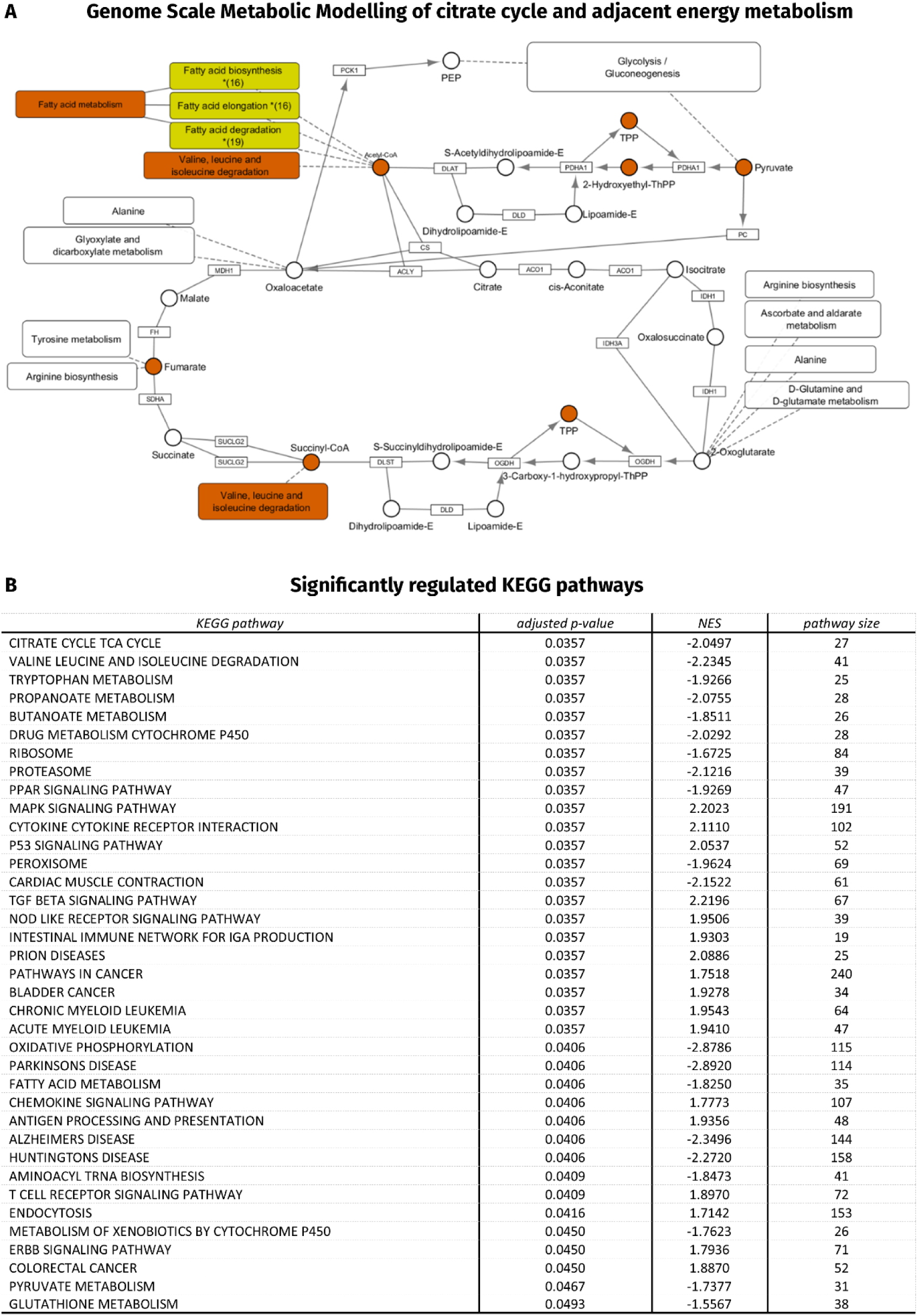
Genome Scale Metabolic Modelling (GEM) and KEGG mapping. (A) KEGG mapping of the results of GEM analysis of endurance athletes (EG) performing acute endurance exercise (EE) comparing post timepoint with pre. Shown is the significantly affected citrate cycle pathway together with neighboring metabolic pathways. Orange color are reporter metabolites and pathways that are significantly downregulated. Yellow color are pathways in which a high number of their member reporter metabolites are downregulated (number of reporter metabolites in parenthesis). (B) Significantly regulated KEGG pathways in endurance trained athletes comparing post with pre timepoint immediately following acute endurance exercise.

**Figure S6:**
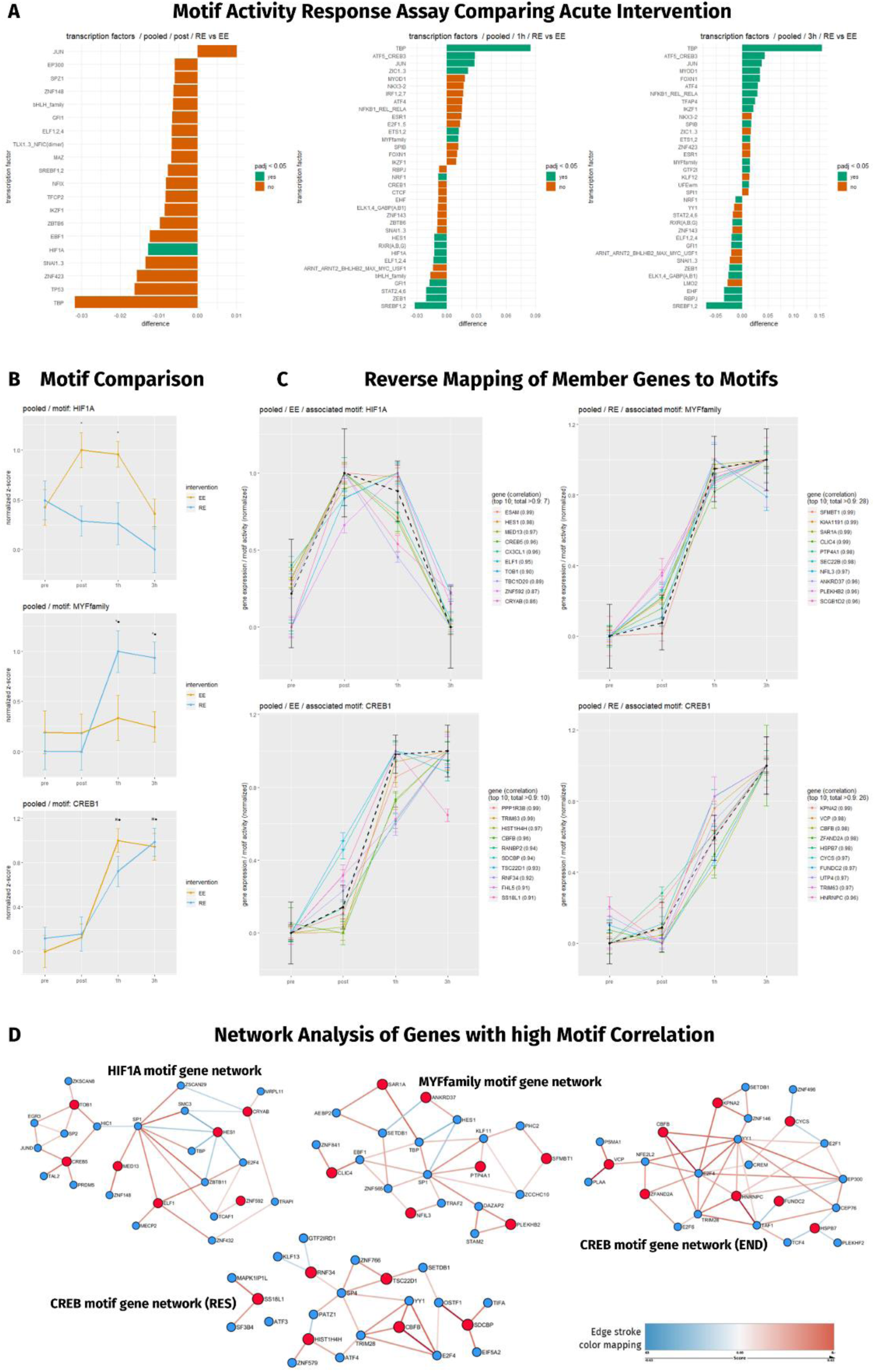
Motif Activity Response Assay (MARA) and network analysis using TCSBN. (A) MARA comparing acute endurance (EE) and resistance exercise (RE) for all groups pooled at post, 1h and 3h timepoints (B) Direct comparison of motif activity time-course for the motifs HIF1A, MYFfamily and CREB. (C) Reverse mapping of motif member genes to motif activity (dashed line) using curve correlation. Top 10 genes are plotted, number of genes with correlation > 0.9 and individual gene correlation in parenthesis. (D) Network analysis of genes identified in reverse mapping (red dots) using the tissue and cancer specific biological networks (TCSBN) database. Edge colors represent positive (red) or negative (blue) connection of genes within the network.

## REFERENCES

1. Lindholm ME, Marabita F, Gomez-Cabrero D, et al. An integrative analysis reveals coordinated reprogramming of the epigenome and the transcriptome in human skeletal muscle after training. Epigenetics. 2014;9(12):1557–1569. doi:10.4161/15592294.2014.982445

2. Stepto NK, Coffey VG, Carey AL, et al. Global gene expression in skeletal muscle from well-trained strength and endurance athletes. Med Sci Sports Exerc. 2009;41(3):546–565. doi:10.1249/MSS.0b013e31818c6be9

3. Seaborne RA, Sharples AP. The interplay between exercise metabolism, epigenetics, and skeletal muscle remodeling. Exerc Sport Sci Rev. 2020;48(4):188–200. doi:10.1249/JES.0000000000000227

4. Perry CGR, Lally J, Holloway GP, Heigenhauser GJF, Bonen A, Spriet LL. Repeated transient mRNA bursts precede increases in transcriptional and mitochondrial proteins during training in human skeletal muscle. J Physiol. 2010;588(23):4795–4810. doi:10.1113/jphysiol.2010.199448

5. Coffey VG, Zhong Z, Shield A, et al. Early signaling responses to divergent exercise stimuli in skeletal muscle from well-trained humans. FASEB J. 2006;20(1):190–192. doi:10.1096/fj.05-4809fje

6. Robinson MM, Dasari S, Konopka AR, et al. Enhanced Protein Translation Underlies Improved Metabolic and Physical Adaptations to Different Exercise Training Modes in Young and Old Humans. Cell Metab. 2017;25(3):581–592. doi:10.1016/j.cmet.2017.02.009

7. Dickinson JM, D’Lugos AC, Naymik MA, et al. Transcriptome response of human skeletal muscle to divergent exercise stimuli. J Appl Physiol. Published online 2018:japplphysiol.00014.2018. doi:10.1152/japplphysiol.00014.2018

8. Rundqvist HC, Montelius A, Osterlund T, et al. Acute sprint exercise transcriptome in human skeletal muscle. PLoS One. 2019;14(10):1–24. doi:10.1371/journal.pone.0223024

9. Contrepois K, Wu S, Moneghetti KJ, et al. Molecular Choreography of Acute Exercise. Cell. 2020;181(5):1112–1130.e16. doi:10.1016/j.cell.2020.04.043

10. Chapman MA, Arif M, Emanuelsson EB, et al. Skeletal Muscle Transcriptomic Comparison between Long-Term Trained and Untrained Men and Women. Cell Rep. 2020;31(12). doi:10.1016/j.celrep.2020.107808

11. Holloszy JO, Oscai LB, Don IJ, Molé PA. Mitochondrial citric acid cycle and related enzymes: Adaptive response to exercise. Biochem Biophys Res Commun. 1970;40(6):1368–1373. doi:https://doi.org/10.1016/0006-291X(70)90017-3

12. Ørtenblad N, Nielsen J, Boushel R, Söderlund K, Saltin B, Holmberg HC. The muscle fiber profiles, mitochondrial content, and enzyme activities of the exceptionally well-trained arm and leg muscles of elite cross-country skiers. Front Physiol. 2018;9(AUG):1–11. doi:10.3389/fphys.2018.01031

13. Tesch PA, Karlsson J. Muscle fiber types and size in trained and untrained muscles of elite athletes. J Appl Physiol. 1985;59(6):1716–1720. doi:10.1152/jappl.1985.59.6.1716

14. Vollaard NBJ, Constantin-Teodosiu D, Fredriksson K, et al. Systematic analysis of adaptations in aerobic capacity and submaximal energy metabolism provides a unique insight into determinants of human aerobic performance. J Appl Physiol. 2009;106(5):1479–1486. doi:10.1152/japplphysiol.91453.2008

15. Wucher V, Sodaei R, Amador R, Irimia M, Guigó R. Day-night and seasonal variation of human gene expression across tissues. bioRxiv. Published online January 1, 2022:2021.02.28.433266. doi:10.1101/2021.02.28.433266

16. Chukwuma CI, Mopuri R, Nagiah S, Chuturgoon AA, Islam MS. Erythritol reduces small intestinal glucose absorption, increases muscle glucose uptake, improves glucose metabolic enzymes activities and increases expression of Glut-4 and IRS-1 in type 2 diabetic rats. Eur J Nutr. 2018;57(7):2431–2444. doi:10.1007/s00394-017-1516-x

17. Sánchez OA, Walseth TF, Snow LM, Serfass RC, Thompson L V. Skeletal muscle sorbitol levels in diabetic rats with and without insulin therapy and endurance exercise training. Exp Diabetes Res. 2009;2009:737686. doi:10.1155/2009/737686

18. Starnes JW, Parry TL, O’Neal SK, et al. Exercise-induced alterations in skeletal muscle, heart, liver, and serum metabolome identified by non-targeted metabolomics analysis. Metabolites. 2017;7(3). doi:10.3390/metabo7030040

19. Nada MA, Abdel-Aleem S, Schulz H. On the rate-limiting step in the β-oxidation of polyunsaturated fatty acids in the heart. Biochim Biophys Acta (BBA)/Lipids Lipid Metab. 1995;1255(3):244–250. doi:10.1016/0005-2760(94)00223-L

20. Foster DW. The role of the carnitine system in human metabolism. Ann N Y Acad Sci. 2004;1033:1–16. doi:10.1196/annals.1320.001

21. Puigarnau S, Fernàndez A, Obis E, et al. Metabolomics reveals that fittest trail runners show a better adaptation of bioenergetic pathways. J Sci Med Sport. 2022;(xxxx). doi:10.1016/j.jsams.2021.12.006

22. Longo N, Frigeni M, Pasquali M. Carnitine transport and fatty acid oxidation. Biochim Biophys Acta - Mol Cell Res. 2016;1863(10):2422–2435. doi:10.1016/j.bbamcr.2016.01.023

23. Saha S, Anilkumar AA, Mayor S. GPI-anchored protein organization and dynamics at the cell surface. J Lipid Res. 2016;57(2):159–175. doi:10.1194/jlr.R062885

24. Nadiv O, Shinitzky M, Manu H, et al. Elevated protein tyrosine phosphatase activity and increased membrane viscosity are associated with impaired activation of the insulin receptor kinase in old rats. Biochem J. 1994;298(2):443–450. doi:10.1042/bj2980443

25. Görski J, Zendzian-Piotrowska M, De Jong YF, Niklińska W, Glatz JFC. Effect of endurance training on the phospholipid content of skeletal muscles in the rat. Eur J Appl Physiol Occup Physiol. 1999;79(5):421–425. doi:10.1007/s004210050532

26. Heden TD, Neufer PD, Funai K. Looking Beyond Structure: Membrane Phospholipids of Skeletal Muscle Mitochondria.

27. Trends Endocrinol Metab. 2016;27(8):553–562. doi:10.1016/j.tem.2016.05.007

27. Lee S, Norheim F, Gulseth HL, et al. Skeletal muscle phosphatidylcholine and phosphatidylethanolamine respond to exercise and influence insulin sensitivity in men. Sci Rep. 2018;8(1):1–12. doi:10.1038/s41598-018-24976-x

28. Ismail AD, Alkhayl FFA, Wilson J, Johnston L, Gill JMR, Gray SR. The effect of short-duration resistance training on insulin sensitivity and muscle adaptations in overweight men. Exp Physiol. 2019;104(4):540–545. doi:10.1113/EP087435

29. Leto D, Saltiel AR. Regulation of glucose transport by insulin: Traffic control of GLUT4. Nat Rev Mol Cell Biol. 2012;13(6):383–396. doi:10.1038/nrm3351

30. Amar D, Lindholm ME, Norrbom J, Wheeler MT, Rivas MA, Ashley EA. Time trajectories in the transcriptomic response to exercise - a meta-analysis. Nat Commun. 2021;12(1):1–12. doi:10.1038/s41467-021-23579-x

31. Freyssenet D, Berthon P, Denis C. Mitochondrial Biogenesis in Skeletal Muscle in Response to Endurance Exercises. Arch Physiol Biochem. 1996;104(2):129–141. doi:10.1076/apab.104.2.129.12878

32. Nielsen J, Gejl KD, Hey-Mogensen M, et al. Plasticity in mitochondrial cristae density allows metabolic capacity modulation in human skeletal muscle. J Physiol. 2017;595(9):2839–2847. doi:10.1113/JP273040

33. Porter C, Reidy PT, Bhattarai N, Sidossis LS, Rasmussen BB. Resistance Exercise Training Alters Mitochondrial Function in Human Skeletal Muscle. Med Sci Sports Exerc. 2015;47(9):1922–1931. doi:10.1249/MSS.0000000000000605

34. Tesch P, Thorsson A, Colliander E. Effects of eccentric and concentric resistance training on skeletal muscle substrates, enzyme activities and capillary supply. Acta Physiol Scand. 1990;140(4):575–579. doi:https://doi.org/10.1111/j.1748-1716.1990.tb09035.x

35. Siu PM, Donley DA, Bryner RW, Alway SE. Citrate synthase expression and enzyme activity after endurance training in cardiac and skeletal muscles. J Appl Physiol. 2003;94(2):555–560. doi:10.1152/japplphysiol.00821.2002

36. Mardinoglu A, Agren R, Kampf C, Asplund A, Uhlen M, Nielsen J. Genome-scale metabolic modelling of hepatocytes reveals serine deficiency in patients with non-alcoholic fatty liver disease. Nat Commun. 2014;5(May 2013):1–11. doi:10.1038/ncomms4083

37. MacQueen J. Some methods for classification and analysis of multivariate observations. In: 1967.

38. Chukwuma CI, Islam S. Sorbitol increases muscle glucose uptake ex vivo and inhibits intestinal glucose absorption ex vivo and in normal and type 2 diabetic rats. Appl Physiol Nutr Metab. 2017;42(4):377–383. doi:10.1139/apnm-2016-0433

39. Hayashi T, Hirshman MF, Fujii N, Habinowski SA, Witters LA, Goodyear LJ. Metabolic stress and altered glucose transport: activation of AMP-activated protein kinase as a unifying coupling mechanism. Diabetes. 2000;49(4):527–531. doi:10.2337/diabetes.49.4.527

40. Borum PR. Carnitine. Annu Rev Nutr. 1983;3(1):233–259. doi:10.1146/annurev.nu.03.070183.001313

41. Longo N, Amat Di San Filippo C, Pasquali M. Disorders of carnitine transport and the carnitine cycle. Am J Med Genet - Semin Med Genet. 2006;142 C(2):77–85. doi:10.1002/ajmg.c.30087

42. Drosatos K, Schulze PC. Cardiac lipotoxicity: Molecular pathways and therapeutic implications. Curr Heart Fail Rep. 2013;10(2):109–121. doi:10.1007/s11897-013-0133-0

43. Forrest ARR, Kawaji H, Rehli M, et al. A promoter-level mammalian expression atlas. Nature. 2014;507(7493):462–470. doi:10.1038/nature13182

44. Arner E, Daub CO, Vitting-Seerup K, et al. Transcribed enhancers lead waves of coordinated transcription in transitioning mammalian cells. Science (80-). 2015;347(6225):1010 LP–1014. doi:10.1126/science.1259418

45. Suzuki H, Forrest ARR, van Nimwegen E, et al. The transcriptional network that controls growth arrest and differentiation in a human myeloid leukemia cell line. Nat Genet. 2009;41(5):553–562. doi:10.1038/ng.375

46. Vogt M, Puntschart A, Geiser J, Zuleger C, Billeter R, Hoppeler H. Molecular adaptations in human skeletal muscle to endurance training under simulated hypoxic conditions. J Appl Physiol. 2001;91(1):173–182. doi:10.1152/jappl.2001.91.1.173

47. Schofield CJ, Ratcliffe PJ. Oxygen sensing by HIF hydroxylases. Nat Rev Mol Cell Biol. 2004;5(5):343–354. doi:10.1038/nrm1366

48. Bigham AW, Lee FS. Human high-altitude adaptation: Forward genetics meets the HIF pathway. Genes Dev. 2014;28(20):2189–2204. doi:10.1101/gad.250167.114

49. Burton DA, Stokes K, Hall GM. Physiological effects of exercise. Contin Educ Anaesth Crit Care Pain. 2004;4(6):185–188. doi:10.1093/bjaceaccp/mkh050

50. Huang LE, Arany Z, Livingston DM, Franklin Bunn H. Activation of hypoxia-inducible transcription factor depends primarily upon redox-sensitive stabilization of its α subunit. J Biol Chem. 1996;271(50):32253–32259. doi:10.1074/jbc.271.50.32253

51. Maxwell PH, Wiesener MS, Chang G-W, et al. The tumour suppressor protein VHL targets hypoxia-inducible factors for oxygen-dependent proteolysis. Nature. 1999;399(20 May):271–275.

52. Wang GL, Jiang BH, Rue EA, Semenza GL. Hypoxia-inducible factor 1 is a basic-helix-loop-helix-PAS heterodimer regulated by cellular O2 tension. Proc Natl Acad Sci U S A. 1995;92(12):5510–5514. doi:10.1073/pnas.92.12.5510

53. Hernández-Hernández JM, García-González EG, Brun CE, Rudnicki MA. The myogenic regulatory factors, determinants of muscle development, cell identity and regeneration. Semin Cell Dev Biol. 2017;72:10–18. doi:10.1016/j.semcdb.2017.11.010

54. Herzig S, Long F, Jhala US, et al. CREB regulates hepatic gluconeogenesis through the coactivator PGC-1. Nature. 2001;413(6852):179–183. doi:10.1038/35093131

55. Handschin C, Rhee J, Lin J, Tarr PT, Spiegelman BM. An autoregulatory loop controls peroxisome proliferator-activated receptor γ coactivator 1α expression in muscle. Proc Natl Acad Sci U S A. 2003;100(12):7111–7116. doi:10.1073/pnas.1232352100

56. Balwierz PJ, Pachkov M, Arnold P, et al. ISMARA: automated modeling of genomic signals as a democracy of regulatory motifs. Genome Res. 2014;24(5):869–884. doi:10.1101/gr.169508.113

57. Noguchi YT, Nakamura M, Hino N, et al. Cell-autonomous and redundant roles of Hey1 and HeyL in muscle stem cells: HeyL requires HeS1 to bind diverse DNA sites. Dev. 2019;146(4):1–12. doi:10.1242/dev.163618

58. Tung JJ, Hobert O, Berryman M, Kitajewski J. Chloride intracellular channel 4 is involved in endothelial proliferation and morphogenesis in vitro. Angiogenesis. 2009;12(3):209–220. doi:10.1007/s10456-009-9139-3

59. Bodine SC, Latres E, Baumhueter S, et al. Identification of ubiquitin ligases required for skeletal Muscle Atrophy. Science (80-). 2001;294(5547):1704–1708. doi:10.1126/science.1065874

60. Philipot O, Joliot V, Ait-Mohamed O, et al. The core binding factor CBF negatively regulates skeletal muscle terminal differentiation. PLoS One. 2010;5(2). doi:10.1371/journal.pone.0009425

61. Schoenfeld BJ. Potential mechanisms for a role of metabolic stress in hypertrophic adaptations to resistance training. Sport Med. 2013;43(3):179–194. doi:10.1007/s40279-013-0017-1

62. Nieman DC, Wentz LM. The compelling link between physical activity and the body’s defense system. J Sport Heal Sci. 2019;8(3):201–217. doi:10.1016/j.jshs.2018.09.009

63. Pillon NJ, Gabriel BM, Dollet L, et al. Transcriptomic profiling of skeletal muscle adaptations to exercise and inactivity. Nat Commun. 2020;11(1). doi:10.1038/s41467-019-13869-w

64. Arany Z, Foo S-Y, Ma Y, et al. HIF-independent regulation of VEGF and angiogenesis by the transcriptional coactivator PGC-1a. Yale J Biol Med. 2008;451:1008–1013. doi:10.1038/nature06613

65. Chinsomboon J, Ruas J, Gupta RK, et al. The transcriptional coactivator PGC-1alpha mediates exercise-induced angiogenesis in skeletal muscle. Proc Natl Acad Sci U S A. 2009;106(50):21401–21406. doi:10.1073/pnas.0909131106

66. Pilegaard H, Saltin B, Neufer PD. Exercise induces transient transcriptional activation of the PGC-1α gene in human skeletal muscle. J Physiol. 2003;546(3):851–858. doi:10.1113/jphysiol.2002.034850

67. Ruas JL, White JP, Rao RR, et al. A PGC-1α isoform induced by resistance training regulates skeletal muscle hypertrophy. Cell. 2012;151(6):1319–1331. doi:10.1016/j.cell.2012.10.050

68. Ydfors M, Fischer H, Mascher H, Blomstrand E, Norrbom J, Gustafsson T. The truncated splice variants, NT-PGC-1α and PGC-1α4, increase with both endurance and resistance exercise in human skeletal muscle. Physiol Rep. 2013;1(6):1–9. doi:10.1002/phy2.140

69. Buckingham M, Rigby PWJJ. Gene Regulatory Networks and Transcriptional Mechanisms that Control Myogenesis. Dev Cell. 2014;28(3):225–238. doi:10.1016/j.devcel.2013.12.020

70. Yang Y, Creer A, Jemiolo B, Trappe S. Time course of myogenic and metabolic gene expression in response to acute exercise in human skeletal muscle. J Appl Physiol. 2005;98(5):1745–1752. doi:10.1152/japplphysiol.01185.2004

71. McHugh MP. Recent advances in the understanding of the repeated bout effect: The protective effect against muscle damage from a single bout of eccentric exercise. Scand J Med Sci Sport. 2003;13(2):88–97. doi:10.1034/j.1600-0838.2003.02477.x

72. Moro C, Bajpeyi S, Smith SR. Determinants of intramyocellular triglyceride turnover: Implications for insulin sensitivity. Am J Physiol - Endocrinol Metab. 2008;294(2). doi:10.1152/ajpendo.00624.2007

73. Scharhag-Rosenberger F, Meyer T, Walitzek S, Kindermann W. Effects of one year aerobic endurance training on resting metabolic rate and exercise fat oxidation in previously untrained men and women. Metabolic endurance training adaptations. Int J Sports Med. 2010;31(7):498–504. doi:10.1055/s-0030-1249621

74. Schenk S, Horowitz JF. Coimmunoprecipitation of FAT/CD36 and CPT I in skeletal muscle increases proportionally with fat oxidation after endurance exercise training. Am J Physiol - Endocrinol Metab. 2006;291(2):254–260. doi:10.1152/ajpendo.00051.2006

75. Jackman SR, Witard OC, Philp A, Wallis GA, Baar K, Tipton KD. Branched-chain amino acid ingestion stimulates muscle myofibrillar protein synthesis following resistance exercise in humans. Front Physiol. 2017;8(JUN). doi:10.3389/fphys.2017.00390

76. Kim D-H, Kim S-H, Jeong W-S, Lee H-Y. Effect of BCAA intake during endurance exercises on fatigue substances, muscle damage substances, and energy metabolism substances. J Exerc Nutr Biochem. 2013;17(4):169–180. doi:10.5717/jenb.2013.17.4.169

77. Sharp CPM, Pearson DR. Amino acid supplements and recovery from high-intensity resistance training. J Strength Cond Res. 2010;24(4):1125–1130. doi:10.1519/JSC.0b013e3181c7c655

78. Weber MG, Dias SS, de Angelis TR, et al. The use of BCAA to decrease delayed-onset muscle soreness after a single bout of exercise: a systematic review and meta-analysis. Amino Acids. 2021;53(11):1663–1678. doi:10.1007/s00726-021-03089-2

79. Yuen M, Cooper ST, Marston SB, et al. Muscle weakness in TPM3-myopathy is due to reduced Ca2+-sensitivity and impaired acto-myosin cross-bridge cycling in slow fibres. Hum Mol Genet. 2015;24(22):6278–6292. doi:10.1093/hmg/ddv334

80. Altomonte J, Cong L, Harbaran S, et al. Foxo1 mediates insulin action on apoC-III and triglyceride metabolism. J Clin Invest. 2004;114(10):1493–1503. doi:10.1172/JCI200419992

81. Puigserver P, Rhee J, Donovan J, et al. Insulin-regulated hepaticgluconeogenesis throughFOXO1–PGC-1ainteractio. Nature. 2003;423(6939):545–550. doi:10.1038/nature01606.1.

82. Samuel VT, Choi CS, Phillips TG, et al. Targeting Foxo1 in mice using antisense oligonucleotide improves hepatic and peripheral insulin action. Diabetes. 2006;55(7):2042–2050. doi:10.2337/db05-0705

83. Matsumoto M, Pocai A, Rossetti L, DePinho RA, Accili D. Impaired Regulation of Hepatic Glucose Production in Mice Lacking the Forkhead Transcription Factor Foxo1 in Liver. Cell Metab. 2007;6(3):208–216. doi:10.1016/j.cmet.2007.08.006

84. Gross DN, Van Den Heuvel APJ, Birnbaum MJ. The role of FoxO in the regulation of metabolism. Oncogene. 2008;27(16):2320–2336. doi:10.1038/onc.2008.25

85. Fyfe JJ, Bishop DJ, Bartlett JD, et al. Enhanced skeletal muscle ribosome biogenesis, yet attenuated mTORC1 and ribosome biogenesis-related signalling, following short-term concurrent versus single-mode resistance training. Sci Rep. 2018;8(1):1–21. doi:10.1038/s41598-017-18887-6

86. Bodine SC, Stitt TN, Gonzalez M, et al. Akt/mTOR pathway is a crucial regulator of skeletal muscle hypertrophy and can prevent muscle atrophy in vivo. Nat Cell Biol. 2001;3(November):1014–1019.

87. Mazo CE, D’Lugos AC, Sweeney KR, et al. The effects of acute aerobic and resistance exercise on mTOR signaling and autophagy markers in untrained human skeletal muscle. Eur J Appl Physiol. 2021;121(10):2913–2924. doi:10.1007/s00421-021-04758-6

88. Winder WW, Hardie DG. AMP-activated protein kinase, a metabolic master switch: possible roles in Type 2 diabetes. Am J Physiol Metab. 1999;277(1):E1–E10. doi:10.1152/ajpendo.1999.277.1.E1

89. Marin TL, Gongol B, Zhang F, et al. AMPK promotes mitochondrial biogenesis and function by phosphorylating the epigenetic factors DNMT1, RBBP7, and HAT1. Sci Signal. 2017;10(464). doi:10.1126/scisignal.aaf7478

90. Ponticos M, Lu QL, Morgan JE, Hardie DG, Partridge TA, Carling D. Dual regulation of the AMP-activated protein kinase provides a novel mechanism for the control of creatine kinase in skeletal muscle. EMBO J. 1998;17(6):1688–1699. doi:10.1093/emboj/17.6.1688

91. Mantovani J, Roy R. Re-evaluating the general(ized) roles of AMPK in cellular metabolism. FEBS Lett. 2011;585(7):967–972. doi:10.1016/j.febslet.2010.12.015

92. Bengal E, Aviram S, Hayek T. P38 mapk in glucose metabolism of skeletal muscle: Beneficial or harmful? Int J Mol Sci. 2020;21(18):1–17. doi:10.3390/ijms21186480

93. Sakai M, Matsumoto M, Tujimura T, et al. CITED2 links hormonal signaling to PGC-1α acetylation in the regulation of gluconeogenesis. Nat Med. 2012;18(4):612–617. doi:10.1038/nm.2691

94. Yoon H, Lim JH, Cho CH, Huang LE, Park JW. CITED2 controls the hypoxic signaling by snatching p300 from the two distinct activation domains of HIF-1α. Biochim Biophys Acta - Mol Cell Res. 2011;1813(12):2008–2016. doi:10.1016/j.bbamcr.2011.08.018

95. Ruiz-Ortiz I, De Sancho D. Competitive binding of HIF-1α and CITED2 to the TAZ1 domain of CBP from molecular simulations. Phys Chem Chem Phys. 2020;22(15):8118–8127. doi:10.1039/d0cp00328j

96. Kolobova E, Tuganova A, Boulatnikov I, Popov KM. Regulation of pyruvate dehydrogenase activity through phosphorylation at multiple sites. Biochem J. 2001;358(1):69–77. doi:10.1042/0264-6021:3580069

97. Luque A, Carpizo DR, Iruela-Arispe ML. ADAMTS1/METH1 inhibits endothelial cell proliferation by direct binding and sequestration of VEGF165. J Biol Chem. 2003;278(26):23656–23665. doi:10.1074/jbc.M212964200

98. Hawley JA. Adaptations of skeletal muscle to prolonged, intense endurance training. Clin Exp Pharmacol Physiol. 2002;29(3):218–222. doi:10.1046/j.1440-1681.2002.03623.x

99. Damas F, Phillips S, Vechin FC, Ugrinowitsch C. A Review of Resistance Training-Induced Changes in Skeletal Muscle Protein Synthesis and Their Contribution to Hypertrophy. Sport Med. 2015;45(6):801–807. doi:10.1007/s40279-015-0320-0

100. Spanidis Y, Stagos D, Papanikolaou C, et al. Resistance-Trained Individuals Are Less Susceptible to Oxidative Damage after Eccentric Exercise. Oxid Med Cell Longev. 2018;2018. doi:10.1155/2018/6857190

101. Moberg M, Lindholm ME, Reitzner SM, Ekblom B, Sundberg C, Psilander N. Exercise Induces Different Molecular Responses in Trained and Untrained Human Muscle. Med Sci Sport Exerc. 2020;(7):1679–1690. doi:10.1249/mss.0000000000002310

102. Hargreaves M, Spriet LL. Skeletal muscle energy metabolism during exercise. Nat Metab. 2020;2(9):817–828. doi:10.1038/s42255-020-0251-4

103. Ekblom-bak E, Engström L-M, Ekblom Ö, Ekblom B. Liv 2000 - Motionsvanor, fysisk prestationsförmåga och levnadsvanor bland svenska kvinnor och män i åldrarna 20-65 år. Published online 2011:1–120.

104. Borg GA. Psychophysical bases of preceived exertion. Med Sci Sport Exerc. 1982;14(5):377–381. doi:-

105. Bergstrom J. Percutaneous needle biopsy of skeletal muscle in physiological and clinical research. Scand J Clin Lab Invest. 1975;35(7):609–616. doi:10.3109/00365517509095787

106. Bass A, Brdiczka D, Eyer P, et al. Metabolic Differentiation of Distinct Muscle Types at the Level of Enzymatic Organization. Eur J Biochem. 1969;10(2):198–206. doi:10.1111/j.1432-1033.1969.tb00674.x

107. A J, Trygg J, Gullberg J, et al. Extraction and GC/MS analysis of the human blood plasma metabolome. Anal Chem. 2005;77(24):8086–8094. doi:10.1021/ac051211v

108. Gullberg J, Jonsson P, Nordström A, Sjöström M, Moritz T. Design of experiments: An efficient strategy to identify factors influencing extraction and derivatization of Arabidopsis thaliana samples in metabolomic studies with gas chromatography/mass spectrometry. Anal Biochem. 2004;331(2):283–295. doi:10.1016/j.ab.2004.04.037

109. Schauer N, Steinhauser D, Strelkov S, et al. GC-MS libraries for the rapid identification of metabolites in complex biological samples. FEBS Lett. 2005;579(6):1332–1337. doi:10.1016/j.febslet.2005.01.029

110. Robinson MD, McCarthy DJ, Smyth GK. edgeR: A Bioconductor package for differential expression analysis of digital gene expression data. Bioinformatics. 2010;26(1):139–140. doi:10.1093/bioinformatics/btp616

111. Benjamini Y, Hochberg Y. Controlling The False Discovery Rate - A Practical And Powerful Approach To Multiple Testing. J R Stat Soc Ser B. 1995;57(1):289–300. doi:10.1111/j.2517-6161.1995.tb02031.x

112. Lee S, Zhang C, Arif M, et al. TCSBN: A database of tissue and cancer specific biological networks. Nucleic Acids Res. 2018;46(D1):D595–D600. doi:10.1093/nar/gkx994

113. Arif M, Zhang C, Li X, et al. INetModels 2.0: An interactive visualization and database of multi-omics data. Nucleic Acids Res. 2021;49(W1):W271–W276. doi:10.1093/nar/gkab254

